# Solving three core challenges in transient dynamics analysis of matrix population models

**DOI:** 10.1101/2025.05.29.656837

**Authors:** Richard A. Hinrichsen

## Abstract

1. Populations are at the mercy of random disturbances large and small and rarely, if ever, converge on predicted long-term behaviours. Therefore, when using matrix population models, ecologists study the dynamics of populations that depart from stable distributions. Necessary for such studies are indices that gauge transient dynamics, which are short-term population fluctuations away from asymptotic trajectories. Conventional indices of transient dynamics present three core challenges: they are distorted by abundant stages of low value (usually immature stages), they are scale dependent, and they conflate transient and asymptotic responses.
2. I develop a new analytical framework for transient dynamics that overcomes these challenges. To solve distortion and scale dependence, I balance a population projection matrix (PPM) by its reproductive values (‘Fisher balancing’) or, alternatively, its stable stage distribution (‘Demetrius balancing’). To disentangle transient and asymptotic dynamics, I strip a PPM of its output in the direction of its stable stage distribution, resulting in a ‘transient’ PPM. I develop indices of transient dynamics that gauge the ‘size’ of a transient PPM using reactivities defined by matrix norms. Among these indices is a new ‘coefficient of transient response’ (COTR) that represents the fraction of the sum of squares of the total response that is explained by the transient response. In addition to the indices of transient dynamics that apply directly to PPMs, the framework also includes case-specific indices of transient dynamics that arise from a specified initial stage vector.
3. Using a dataset of 6,332 PPMs retrieved from the COMPADRE Plant Matrix Database, I compared the new analytical framework with the conventional framework without balancing, focusing mainly on transient responses estimated by COTR. The new framework dramatically changed the rankings of PPMs from the least to the greatest transient dynamics. Spearman correlation between COTR of standardised PPMs and that of balanced PPMs was weak (≤ 0.25). This change in rankings led to new inferences about which taxonomic orders were least or most prone to transient dynamics.
4. By solving three core challenges, the new framework produces a clearer and more robust portrait of transient dynamics.

## 1. INTRODUCTION

Population projection matrix (PPM) models are popular for modelling structured populations because they have a long history of use, use demographic data efficiently, are tractable, and are simple to construct and analyse. They can also answer a host of questions about the effect of increasing or decreasing vital rates with management actions in sensitivity analyses (Caswell, 2001). The classical way to analyse a PPM model is to study its asymptotic dynamics, with an emphasis on stable population growth rate (see, e.g., Otway et al., 2004). In recent decades, however, analysis of transient dynamics has taken centre stage (Stott et al., 2011).

Transient dynamics are the short-term fluctuations of a population away from asymptotic trajectories. A population may deviate far from an asymptotic trajectory because it exists in a dynamic environment subject to perturbations that disrupt population structure. These perturbations can arise from biotic factors such as disease, predation, competition; abiotic factors, such as climate change, extreme weather events, or volcanic eruptions; and anthropogenic factors such as harvest, habitat degradation, or management.

Transient dynamics have implications for population management, comparative demography, and invasive species. Management of renewable resources requires an understanding of transient responses rather than asymptotic behaviour of populations (Hastings et al., 2018). For example, the harvest of bark and foliage of mahogany (*Khaya senegalensis*) in West Africa has a stronger influence on transient dynamics than on asymptotic dynamics (Gaoue, 2016). Population management influences transient dynamics by introducing perturbations that move populations away from asymptotic trajectories (Ezard et al., 2010; Stott et al., 2011). Comparative studies of transient dynamics can guide management actions (Koons et al., 2005; Stott et al., 2010). Monocarpic herbs and trees are more susceptible to transient fluctuations than perennial herbs and shrubs, suggesting different management approaches for these plants (Stott et al., 2010). Lastly, the study of transients helps us understand the dynamics of invasive species (Iles et al., 2016; Stott et al., 2011). Vital rates in young stages of invading plants influence transient dynamics that promote establishment and population growth (Ezard et al., 2010; McMahon & Metcalf, 2008).

Needed for studies of transient dynamics are indices that gauge the size of transient responses, and several such indices are available (Stott et al., 2011). These indices appear in the form of transient bounds that do not require an initial stage vector and case-specific measures that do. Indices of transient dynamics, whether transient bounds or case-specific, exhibit three core challenges: (1) outsized stages with relatively low value have undue influence, (2) they are scale dependent, and (3) they do not separate transient from asymptotic dynamics. These challenge the usefulness of routine inferences made about transient dynamics.

This article aims to present a new analytical framework designed to address these three core challenges. Balancing solves the first two challenges by rescaling the model by reproductive values or by the stable stage distribution (Hinrichsen, 2024). By producing scale-independent indices of transient dynamics, balancing guarantees that, regardless of how stages are counted, as long as alternative counts are rescaled versions of each other (e.g., pre-reproductive or post-reproductive census), indices of transient dynamics are unchanged. To disentangle the transient from asymptotic dynamics, I use stripping, which annihilates the PPM’s output in the direction of the stable stage distribution. The resulting matrix is called a ‘transient’ PPM, which encapsulates the transient response of the original model. Then, I apply indices of transient dynamics directly to the transient PPM. To illustrate this new framework and compare it to a conventional framework, I apply both frameworks to 6,332 matrices selected from the COMPADRE Plant Matrix Database (Salguero-Gómez et al., 2015).

## 2. MATERIALS AND METHODS

### 2.1. Population model

The dynamics of a population are assumed to follow a PPM model of the form

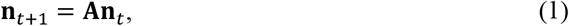

where **n**_*t*_ is the stage vector at time *t* with *n* components, and **A** is an *n* × *n* PPM (Caswell, 2001). This model is linear and discrete with a PPM **A** that is time invariant. The *n* components of the stage vector are abundances of stages, which can represent developmental stages, age classes, size classes, or any combination of these. I assume that PPM **A** is nonnegative with a positive dominant eigenvalue, λ_1_, known as the stable growth rate, and a positive associated right eigenvector, **w**, known as the stable stage distribution. As is customary, I assume that the components of **w** sum to one, and that the reproductive values, which are components of the positive dominant left eigenvector, **v**, are scaled so that **v**^⊤^**w** = 1.

To help disentangle asymptotic from transient dynamics, the standardised PPM, 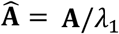, is used instead of **A** (Stott et al., 2011; Stott et al., 2012). The matrix 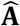 is nonnegative with a dominant eigenvalue of 1 and stable population distribution **w** since

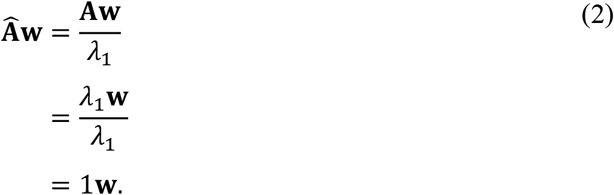

I also use standardised PPMs, but, as we shall see, standardisation is not sufficient to separate transient from asymptotic dynamics.

### 2.2. Indices of transient dynamics

There are several indices that measure the size of transient dynamics of populations (Stott et al., 2011). These depend largely on whether the index is a transient bound or case-specific, and on the time frame used for the measurement. I analyse both transient bounds and case-specific indices over a single time step, highlight their challenges, and propose solutions.

A conventional index of transient dynamics is the reactivity of 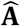. Reactivity is a matrix norm, which is used to gauge the ‘size’ of a matrix. The reactivity of 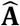 is

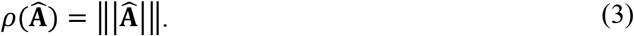

I allow any matrix norm ‖|·|‖ in the definition of reactivity. Some common matrix norms are listed in Table 1 (Horn & Johnson, 2013).

**Table 1.**
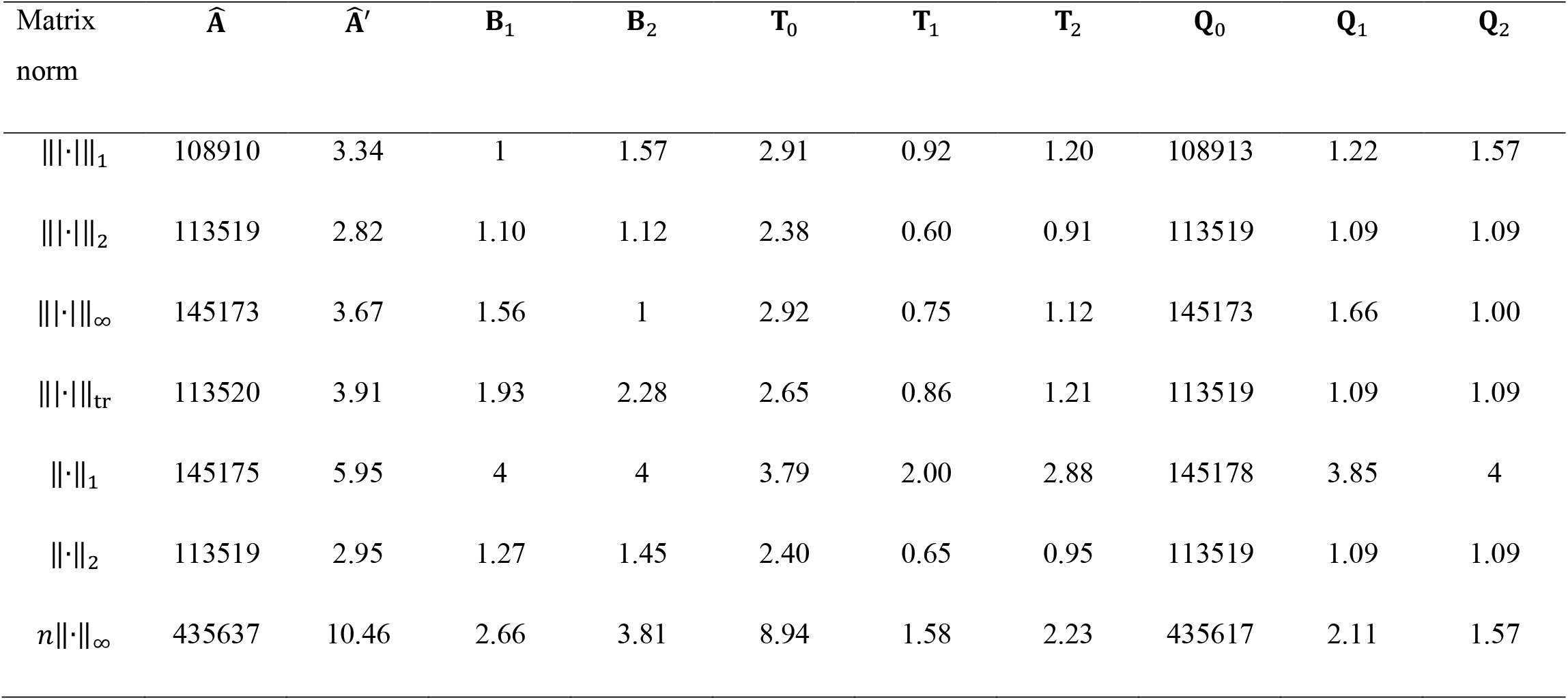
Reactivities for the scarlet monkey flower example. The matrix norms used to define reactivities are the maximum absolute column sum, ‖|·|‖_1_; the spectral norm, ‖|·|‖_2_; and the maximum absolute row sum norm, ‖|·|‖_∞_, the trace norm, ‖|·|‖_tr_, the absolute sum norm, ‖·‖_1_, the Frobenius norm, ‖·‖_2_, and the maximum entry norm, *n*‖·‖_∞_.

The new framework also allows for case-specific reactivities that measure dynamics that arise from a specified initial nonzero stage vector, **n**_0_. Case-specific reactivities are

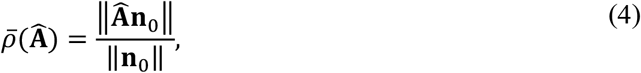

where ‖·‖ is any vector norm. Probably the most common vector norm is the Euclidean norm, ‖·‖_2_, but others are possible (Horn & Johnson, 2013).

The conventional approach of using 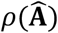 and 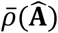 as direct indicators of transient dynamics is problematic, as I will illustrate in the following section. The new framework uses reactivities across a wider range of PPMs, including balanced and transient PPMs, which are derived directly from 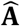.

### 2.3. Core challenges of indices of transient dynamics and their solutions

#### 2.3.1 Undue influence of immature stages

The first challenge of the conventional indices of transient dynamics is that they give undue weight to stages, usually immature stages, that are naturally abundant, but contain individuals of relatively low value. Take, for example, the PPM of the scarlet monkey flower *Mimulus cardinalis* (matrix ID=244375) in COMPADRE. This PPM is

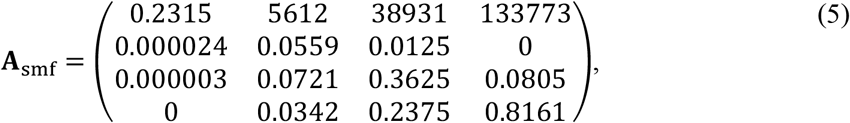

which has 4 stages: (1) seeds, (2) small nonreproductive plants, (3) large nonreproductive plants, and (4) reproductive adults (Angert, 2006). The PPM **A** has stable growth rate 1.23 and stable stage distribution

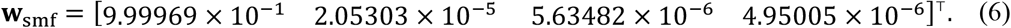

At stability, the seed stage is 200,000 times the number of reproductive adults. Although these components of **w**_smf_ are accurate, they can deceive because not all stages have equal value. Each seed is worth a fraction of the value of a reproductive adult because a seed cannot reproduce and has a very low probability of surviving to maturity. A reproductive adult produces more than 130,000 seeds in one year, and so is worth much more than a single seed. Yet a seed is counted the same as a reproductive adult in conventional indices of transient dynamics. Therefore, seeds exert undue influence on indices. For example, using the Frobenius norm, the reactivity of the scarlet monkey flower standardised PPM is 113519 (Table 1). Reactivities based on the other five norms are also greater than 100,000. These reactivities are distorted by naturally abundant seeds of low value. To solve this challenge, one weights the stages according to their values. For example, one can weight the stages by their reproductive values, which represent expected individual contributions to future generations (Caswell, 2001; Fisher, 1930). For the scarlet monkey flower example, the reproductive values are

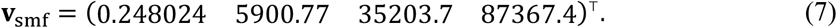

Indeed, according to the reproductive values, a reproductive adult (last stage) is worth much more than a seed (first stage). Weighting stable stage densities by their reproductive values yields

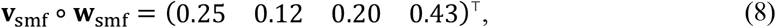

where ∘ denotes a Hadamard ‘component-wise’ product (Horn & Johnson, 2013). Equation (8) gives a fair and undistorted account of the stable stage distribution, with each stage expressed as a fraction of the total reproductive value. Once reproductive values are accounted for, the seed stage is 25% of the stable distribution instead of 99.9969 %. This valuation gives stages a common currency, which allows us to compare stages in a biologically meaningful way. Rescaling stages by their reproductive values is analogous to weighting a collection of pennies, nickels, dimes, and quarters by their respective values of $0.01, $0.05, $0.10, and $0.25. Without such a valuation, it is impossible to know the worth of the coin collection, let alone how much each denomination contributes to that worth.

#### 2.3.2 Distortion

As a measure of the distance between the original and rescaled stable stage distributions, which I call distortion, I use Keyfitz Δ (Caswell, 2001). Using reproductive values to value stages, distortion is

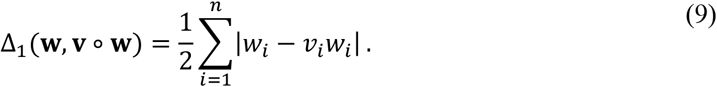

Because Keyfitz Δ is bounded between 0 and 1, it is easy to determine if the distortion is low (near zero) or high (near 1). For the scarlet monkey flower example, the distortion is Δ_1_ = 0.75.

As an alternative to rescaling with reproductive values, one can rescale with the stable stage distribution, where each stage is divided by its corresponding stable stage density (Hinrichsen, 2024). The corresponding alternative measure of distortion is

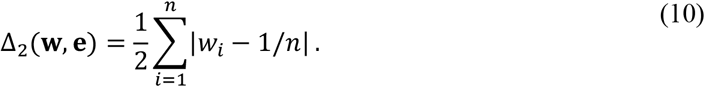

For the monkey flower example, this alternative form of distortion is Δ_2_ = 0.75. Although, in this example, Δ_1_ = Δ_2_, equality of these measures of distortion does not hold in general, and, in fact, Δ_1_ and Δ_2_ can differ greatly for the same PPM.

#### 2.3.3. Scale dependence

Next, I use the scarlet monkey flower example to show that the indices of transient dynamics are scale dependent. Suppose that a plant ecologist, instead of counting seeds individually, counts seeds as the numbers of groups of size 1/*a*_21_, where *a*_21_ is the probability that a seed transitions to the second stage (small nonreproductive stage) in a single year. This new way of counting seeds results in a change of variables and a new PPM, where the new stages are described by

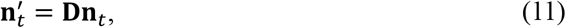

where **D** = diag{(*a*_21_ 1 1 1)^⊤^}. The PPM governing the dynamics of this rescaled stage vector is

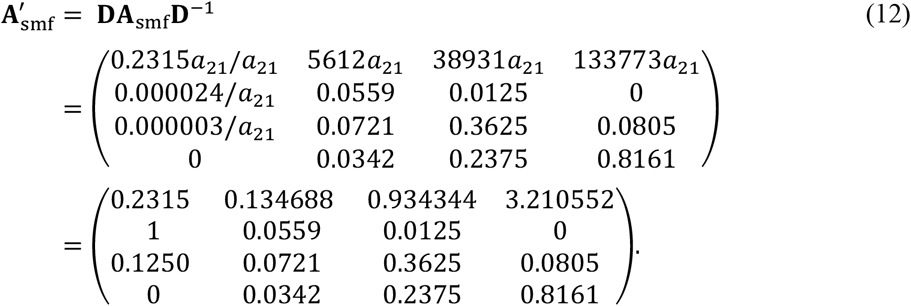

The new PPM 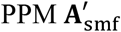 has the same eigenvalues as the original PPM because 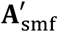 is similar to **A**_smf_ via **D**^−1^ (Horn & Johnson, 2013). In contrast to eigenvalues, which are scale invariant, reactivity differs between the original and rescaled models. For the standardised rescaled PPM, the reactivity using the Frobenius norm is 2.95 (Table 1). Reactivities based on other matrix norms range from 2.82 to 10.46. Compare these reactivities to those of the standardised original PPM, which all exceeded 100,000. Rescaling reduced the reactivities by several orders of magnitude. Reactivities change between the original and rescaled PPMs because they are scale dependent, regardless of the norm employed. It is alarming that reactivities change so much with a simple rescaling of the scarlet monkey flower model. Afterall, the models are fundamentally the same and describe the same population: They are simply rescaled versions of each other.

Scale dependence matters because it can bias a comparative study of transient dynamics, leading to erroneous conclusions. Consider a comparative study in which two identical populations are analysed, except one is counted using a pre-reproductive census and the other, a post-reproductive census (Caswell, 2001; Vindenes et al., 2021). Due to scale dependence, these two census types can produce markedly different values of the indices of transient dynamics, even though the two populations are identical. This challenge can be solved by employing scale-invariant indices of transient dynamics, which are unaffected by the choice of census type. One constructs scale-invariant indices by balancing the PPMs.

#### 2.3.4 Balancing

To prevent the undue influence of certain stages on the indices of transient dynamics, one weights each stage by its value, as in the definitions of distortions. Again, there are two valuations utilized, the first rescales stages by their reproductive values, and the second rescales stages by their stable stage distribution. Rescaling, which is prevalent in mathematical biology, gives ‘small’ and ‘large’ a precise meaning and simplifies a model by reducing the parameters to groupings (Kot, 2001; Murray, 1989; Segel, 1972).

Rescaling stages by reproductive values as in Equation (8) yields

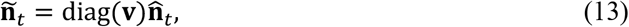

where diag(·) is the square diagonal matrix with diagonal entries equal to the components of the indicated vector, ñ _*t*_ is the rescaled stage vector, and 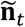 is the stage vector for the standardised matrix model.

Reproductive values are useful for rescaling because they represent individual contributions to future generations, and they distinguish between immature, mature, and post-reproductive individuals (Caswell, 2001). Reproduction, survival, and timing are all included in the reproductive value, which Fisher (1930) identified as the present value of future offspring. Furthermore, the direct action of natural selection is proportional to reproductive value (Fisher, 1930). Fisher (1930) also weighted stages (age classes) by their reproductive values and argued that the weighted numbers gave a more realistic gauge of population growth than the raw unweighted numbers.

When stages are weighted by their reproductive values, each new stage represents the total reproductive value of that stage. The dynamics of the rescaled stage vector ñ_*t*_ are described by the following matrix population model:

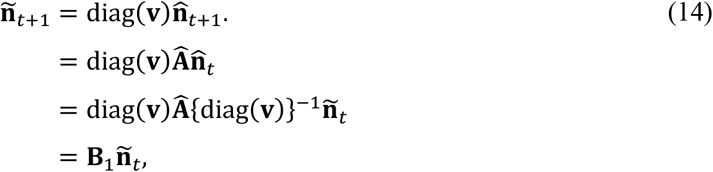

Where

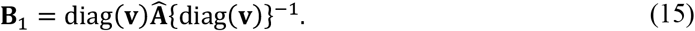

I call **B**_1_ a balanced matrix, and it is the PPM that projects forward the rescaled stage vector ñ _*t*_. **B**_1_ is similar to 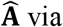 via {diag(**v**)}^−1^, and therefore has the same eigenvalues as 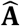 (Horn & Johnson, 2013). Using a diagonal similarity transformation is called balancing in numerical analysis (Golub & Van Loan, 2013), and it is the term I adopt here. Because Fisher (1930) was the first to weight stages by their reproductive values, I call balancing with reproductive values ‘Fisher balancing’.

The vector **e** = (1 1 … 1)^⊤^ is always a left eigenvector of **B**_1_ associated with dominant eigenvalue of 1 since

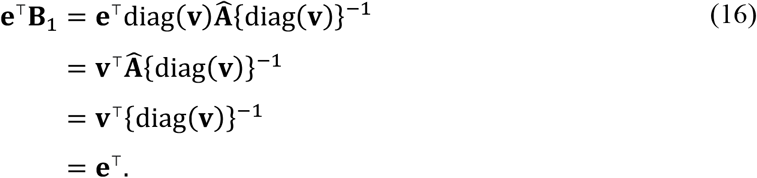

Equation (16) guarantees that each column of **B**_1_ sums to one (i.e. **B**_1_ is column stochastic). The dominant right eigenvector of **B**_1_, also known as the stable stage distribution of **B**_1_, is **v** ∘ **w**. The distortion Δ of the balanced matrix **B** is zero because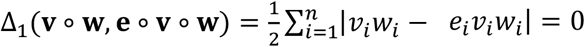.

As an alternative rescaling method, Hinrichsen (2024) rescales stages by the stable stage distribution, yielding

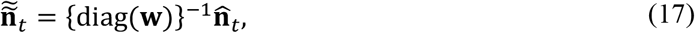

where, 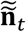 is the rescaled stage vector, and, as before, 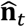 is the stage vector for the standardised matrix model. The matrix that governs the dynamics of the rescaled stage vector is

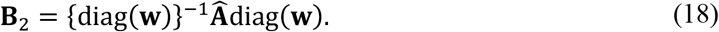

**B**_2_ is an alternative balanced matrix, and it is the PPM that projects forward the rescaled stage vector 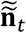. **B**_2_ is similar to 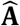 via {diag(**v**)}^−1^, and therefore has the same eigenvalues as 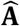 (Horn & Johnson, 2013). The vector **e** = (1 1 … 1)^⊤^ is a right eigenvector of **B**_2_ associated with the dominant eigenvalue of 1. Therefore, each row of **B**_2_ sums to 1 (i.e. **B**_2_ is row stochastic). The stable stage distribution of **B**_2_ is **e**/n, where *n* is the dimensionality of **B**_2_. The reproductive values of **B**_2_ are *n***v** ∘ **w**, which may be verified by multiplying **B**_2_ in Equation (18) on the left by *n*(**v** ∘ **w**)^⊤^. The distortion Δ_2_ of the balanced matrix **B**_2_ is zero because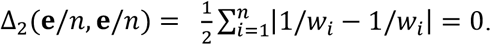.

Rescaling by the stable stage distribution allows us to measure stages in quantities of stable stage density. The procedure gives less value to individuals in naturally abundant stages, such as seeds, and greater value to individuals in naturally less abundant stages, such as reproductive adults. I call balancing with the stable stage distribution ‘Demetrius balancing’ because Demetrius (1974) used the stable stage distribution to rescale a standardised Leslie model. This rescaling produced a balanced Leslie matrix, which he used to derive a measure of entropy.

Balancing ensures that no stage has undue influence on the indices of transient dynamics, because all stages are weighted by their values. A balanced matrix itself is scale invariant and therefore any statistic applied to a balanced PPM, including indices of transient dynamics, will automatically be scale invariant (Appendix S1).

For the scarlet monkey flower example, balancing with reproductive values yields

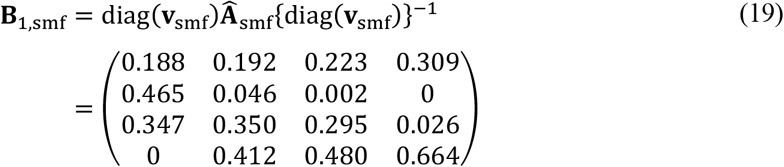

Balancing with the stable stage distribution yields

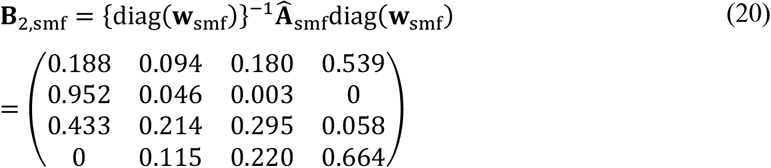

The large differences in the reactivities between the standardised and balanced versions of the scarlet monkey flower matrix are shown in Table 1. Balancing has a profound influence on the ‘size’ of a matrix. When using the Frobenius norm, the reactivities for **B**_1,smf_ and **B**_2,smf_ are 1.27 and 1.45, respectively (Table 1). Reactivities of balanced PPMs using other norms are similarly small, ranging from 1 to 4. Compare these to the reactivities of the standardised original PPM, which all exceeded 100,000. The undue influence of the abundant seed stage on the reactivity of 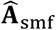 stems from the large fertility rates in the first row, which are not balanced transitions.

#### 2.3.5 Stripping asymptotic components

Balancing remedies the challenges of undue influence of immature stages and scale dependence, but not the final challenge, which requires disentangling transient and asymptotic dynamics (Stott et al., 2011). The output of PPMs, whether balanced or standardised, is not free of the influence of asymptotic dynamics because it contains a component in the direction of the stable stage distribution. This final challenge is solved by a process that I call stripping, whereby one subtracts from each column of a PPM (balanced or not) its component in the direction of the stable stage distribution. Stripping annihilates the asymptotic component of the *output* of a PPM. Haridas and Tuljapurkar (2007) used a similar technique to separate transient and asymptotic components of the *inputs* of PPMs.

Stripping the original 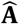 yields a new matrix

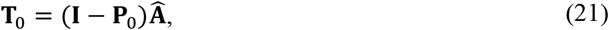

where **I** is the identity matrix and

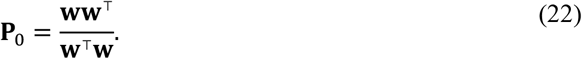

Any component of the output of **T**_0_ in the direction of **w** is annihilated because the columns of **T**_0_ are orthogonal to **w**. Thus, for any input stage vector **n**, its associated output, **T**_0_**n**, represents transient growth alone. This procedure is also applied to balanced matrices.

Stripping **B**_1_ yields the matrix

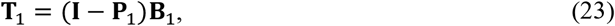

Where

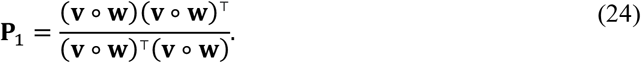

Stripping **B**_2_ yields the matrix

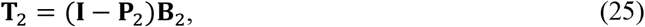

Where

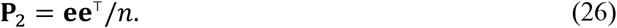

I refer to **T**_0_, **T**_1_, and **T**_2_ as ‘transient’ PPMs because they encapsulate the information needed to infer transient dynamics. I call matrices 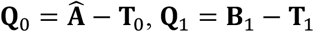, and **Q**_2_ = **B**_2_ − **T**_2_, ‘asymptotic’ PPMs because they encapsulate the information needed to infer asymptotic dynamics.

For the scarlet monkey flower example,

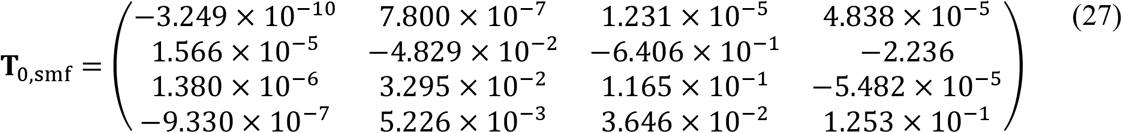

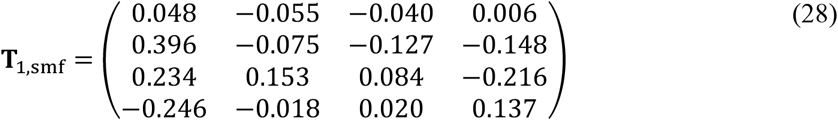

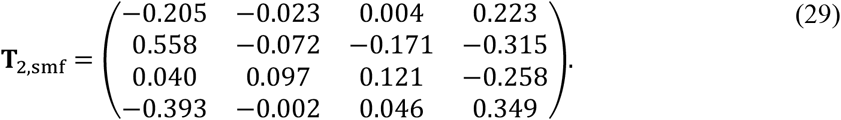

Here, I summarize key properties of transient matrices. Let the eigenvalues of **A** be denoted by λ_1_, λ_2_, …, λ_*n*_. The eigenvalues of the standardised PPM and the balanced PPMs are 1, λ_2_ /λ_1_, …, λ_*n*_ /λ_1_. In contrast, the eigenvalues of the transient matrices are 0, λ_2_ /λ_1_, …, λ_*n*_ / λ_1_. The spectral radius of a transient PPM is given by |λ_2_ |/λ_1_, which is the reciprocal of the damping ratio (Caswell, 2001). A stable stage distribution of a PPM (whether standardised or balanced) is a right eigenvector of its corresponding transient PPM; however, it corresponds to a zero eigenvalue rather than its dominant eigenvalue. This follows because the transient PPM annihilates any stage vector in the direction of the stable stage distribution.

#### 2.3.6 Reactivities of transient matrices

Armed with transient PPMs, we are now ready to calculate meaningful indices of transient dynamics. To do this, one simply calculates the reactivity of the transient PPMs:

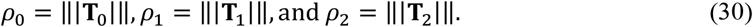

Using the scarlet monkey tree example and defining reactivity as the Frobenius norm, the indices are *ρ*_0_ = 2.40, *ρ*_1_ = 0.65 and *ρ*_2_ = 0.95 (Table 1).

Case-specific indices of transient dynamics are also available using transient matrices, namely,

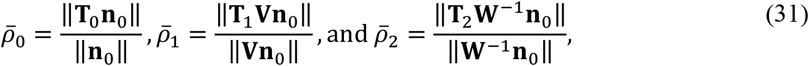

where ‖·‖ represents any vector norm and **n**_0_ is any initial nonzero stage vector. For the scarlet monkey flower example, when **n**_0_ = (1 0 0 0)^⊤^ and the Euclidean norm is used, 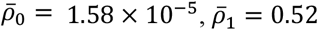, and 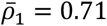.

Reactivities and case-specific reactivities that are based on balanced matrices are scale invariant (Appendix S1). In contrast, because they are derived without balancing, *ρ*_0_ and 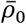 can be badly distorted by the immature stages and they are scale dependent.

#### 2.3.7 Bounds on reactivities

It is helpful to know the bounds on the reactivities of the balanced PPMs and their associated transient PPMs, which I derive in Appendix S2 when reactivity is the Frobenius norm.

Determining which PPMs yield lower and upper bounds gives us insight into what life history characteristics foster small and large transient dynamics. Balancing limits the reactivities of PPMs, providing well-defined and consistent upper bounds. Upper bounds are not presented for the reactivities of PPMs derived without balancing, namely, 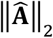, ‖**T**_0_‖_2_, and ‖**Q**_0_‖_2_. Although the lower bounds will still apply, these reactivities can far exceed the upper bounds presented here.

The bounds on the reactivities of balanced PPMs are given by

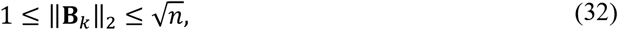

where *k* = 1,2. The lower bound of 1 in Equation (32) also applies to 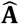, but the upper bound does not: reactivity of 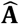 can far exceed 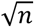, for example 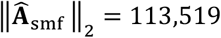 (Table 1).

When *n* > 1, the bounds on the reactivity of transient PPMs **T**_1_ and **T**_2_ are

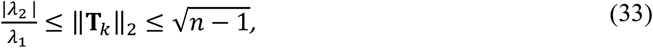

where *k* = 1,2. When *n* = 1, ‖**T**_1_‖_2_ = ‖**T**_2_‖_2_ = 0. When *n* > 1, regardless of the matrix norm used in the definition of reactivity, the reciprocal of the damping ratio is a lower bound on the reactivity of a transient matrix (including **T**_0_) and therefore measured transient dynamics can be zero only if λ_2_ = 0. The upper bound on the reactivity of the transient matrix **T**_0_ can exceed 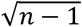, as demonstrated by the scarlet monkey flower example, where ‖**T**_0_‖_2_ = 2.4 (Table 1).

For any *n* ≥ 1, bounds on the reactivities of asymptotic PPMs **Q**_1_ and **Q**_2_ are

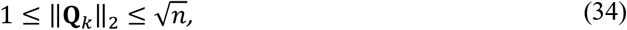

where *k* = 1,2. The upper bound on the reactivity of the transient matrix **Q**_0_ can exceed 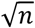, as demonstrated by the scarlet monkey flower example, where ‖**Q**_0_‖_2_ = 113,519 (Table 1).

#### 2.3.8 Coefficient of transient response (COTR)

When using the Frobenius norm, we can take advantage of the fact that the columns of the transient matrix (**T**_*k*_) are orthogonal to those of the corresponding asymptotic matrix (**Q**_*k*_) for *k* = 0,1,2. Orthogonality yields the following identities: 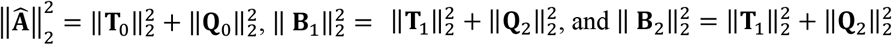 (Appendix S3). Each of these equations decomposes the total sum of squares into transient sum of squares + asymptotic sum of squares. Using these decompositions, I define new indices of transient dynamics:

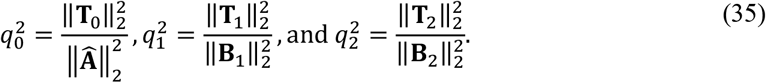

I call each 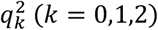 a coefficient of transient response (COTR), which is analogous to the coefficient of determination (i.e. R-squared) from linear regression. The COTR represents the fraction of the total sum of squares that is ‘explained’ by the transient response. Each COTR is bounded as follows: 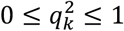 (Appendix S3). When 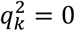, the response is solely asymptotic. When 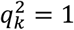, the response is solely transient, which can only be approximately true since there is always some measure of asymptotic response, because 1 ≤ ‖**Q**_*k*_‖_2_ (Appendix S2).

Using the scarlet monkey flower example, the COTRs are 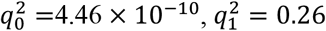, and 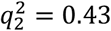.

Case-specific COTRs are also available. Because transient and asymptotic PPMs are orthogonal to one another,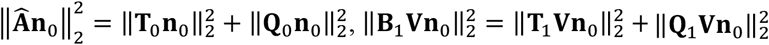, and 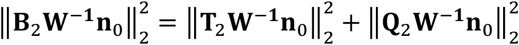 (Appendix S3). This suggests case-specific coefficients that measure the transient response to **n**_0_ as follows:

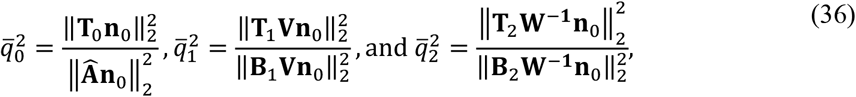

where ‖·‖_2_ is the Euclidean vector norm and **n**_0_ is any initial nonzero stage vector used in the original matrix population model of Equation (1). Case-specific COTRs are bounded as follows: 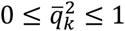 for *k* = 0,1,2 (Appendix S3). When 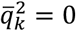, the response is purely asymptotic. When 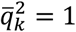 the response is purely transient. A purely transient response is only approximately possible since there will always be a component of **n**_0_ in the direction of the stable stage distribution, **w**. For the scarlet monkey flower example, when 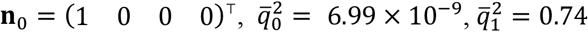, and 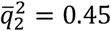.

When they are based on balanced matrices, the COTRs and case-specific COTRs solve all three challenges of conventional indices of transient dynamics: they are not distorted by abundant stages, they are scale invariant (Appendix S1), and they separate transient from asymptotic dynamics. The COTRs 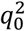 and 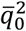 should be used with caution. They allow undue influence of abundant stages of low value and are not scale dependent.

### 2.4. Comparative plant demography application

#### 2.4.1 Dataset of PPMs

For a broad comparison of indices of transient dynamics applied to transient versus standardised PPMs, I retrieved PPMs from the COMPADRE Plant Matrix Database (accessed 5 May 2025, version 6.23.5.0). I accessed COMPADRE via ‘Rcompadre’ (Jones et al., 2022), a package available in the R language and environment for statistical computing and graphics (version 4.5.0, R Core Team, 2025). From COMPADRE, I selected PPMs that contained no missing entries, had dominant eigenvalues that were positive and simple, and were irreducible. To facilitate comparisons among taxonomic orders, I excluded taxonomic orders with fewer than 20 PPMs in COMPADRE. This selection resulted in 6,332 PPMs for populations drawn from 595 species and 34 taxonomic orders. A total of 39% of the PPMs were associated with four taxonomic orders: Asterales, Caryophyllales, Fabales, and Ericales (Figure 1).

**FIGURE 1.**
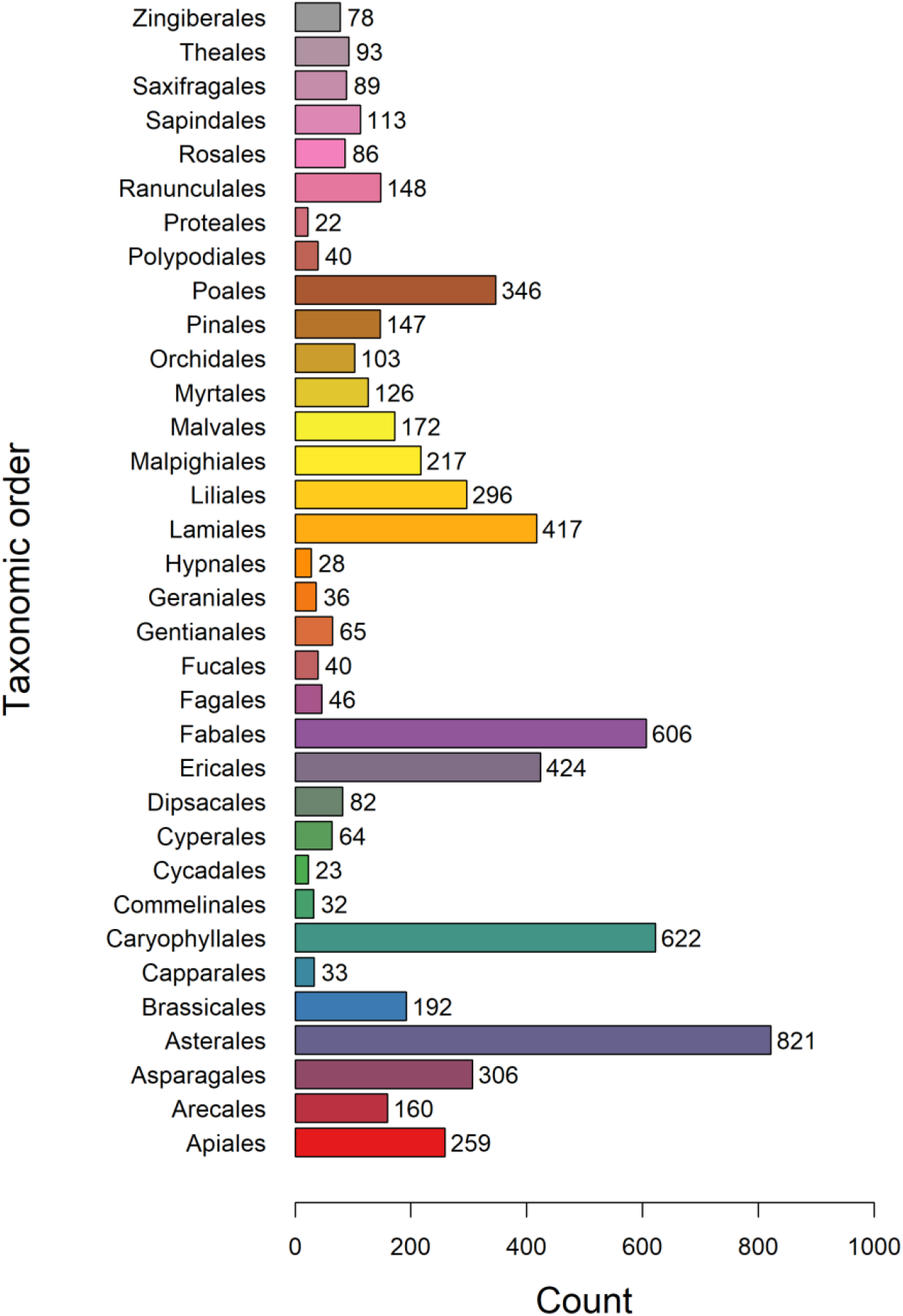
The numbers of PPMs selected from the COMPADRE database by taxonomic order.

The first focus of this application was distortion, where I compared distortions measured using both Δ_1_ and Δ_2_, and described what characteristics of a PPM contribute to high distortions. Next, I compared the indices of transient dynamics, revealing how the balancing method, stripping, distortion, maximum entry, and dimensionality shaped the indices. Throughout the application, I used the Frobenius norm as the measure of reactivity. I compared the new framework with a modified form of the conventional framework, where its transient response is measured as the COTR of the standardized PPM, not its reactivity. This modification avoids confusing transient response with total response.

## 3. RESULTS

### 3.1. Distortions and the role of maximum vital rate

The distribution of distortions Δ_1_ was right skewed, while that of distortions Δ_2_ was bell-shaped with fewer values near the extremes (Figure 2). Despite these distributional differences, there was a tendency for high maximum vital rates to increase distortion, whether measured by Δ_1_ or Δ_2_. Spearman correlations between the maximum vital rate of the standardised PPMs and Δ_1_ and Δ_2_ were strong: *r*_*s*_ = 0.80 and *r*_*s*_ = 0.70, respectively (Figure 3).

**FIGURE 2.**
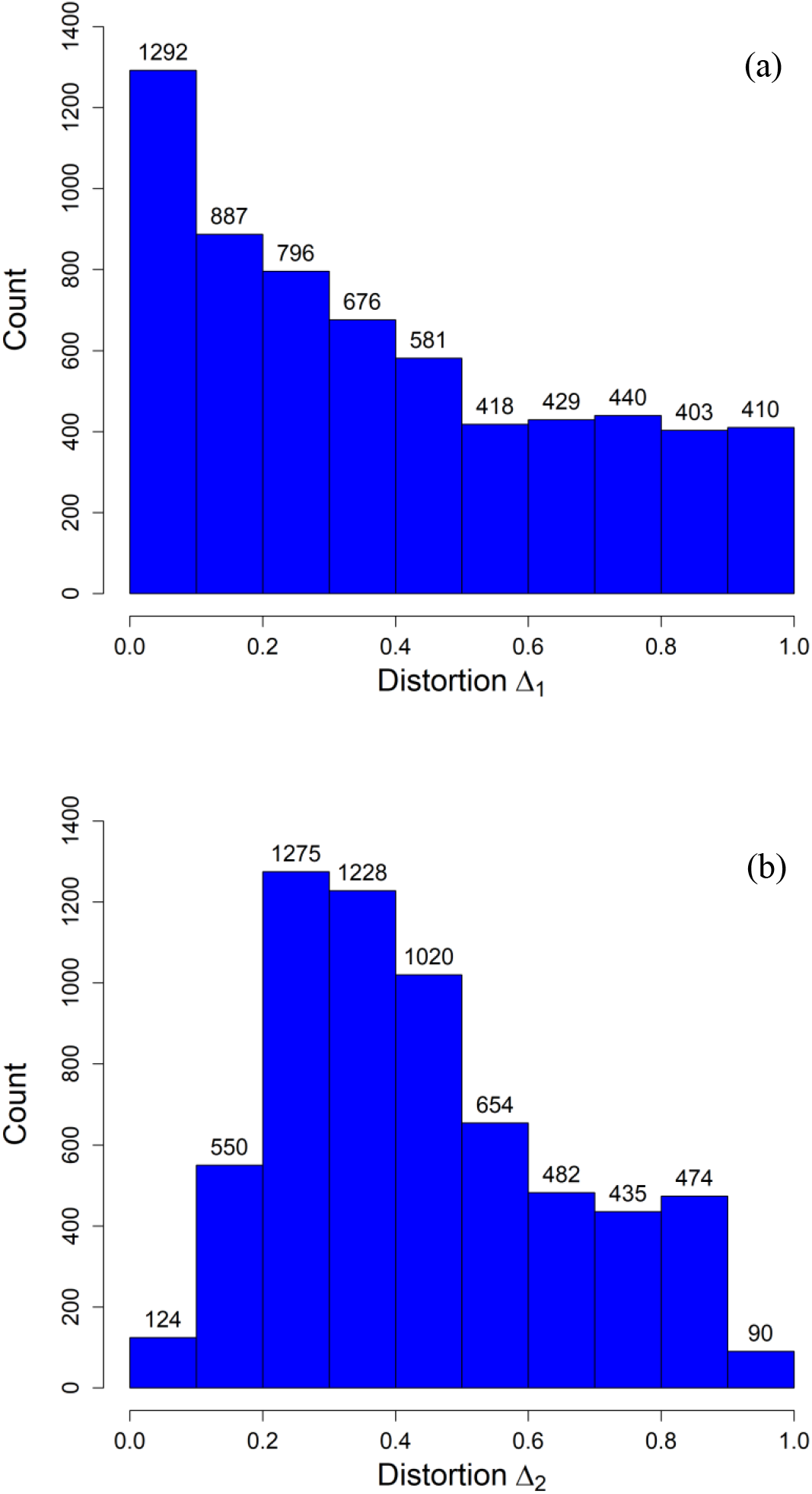
The distribution of distortions of PPMs drawn from the COMPADRE database depicted as histograms for (a) Δ_1_, which is used when rescaling by reproductive values; and (b) Δ_2_, which is used when rescaling by stable stage distribution.

**FIGURE 3.**
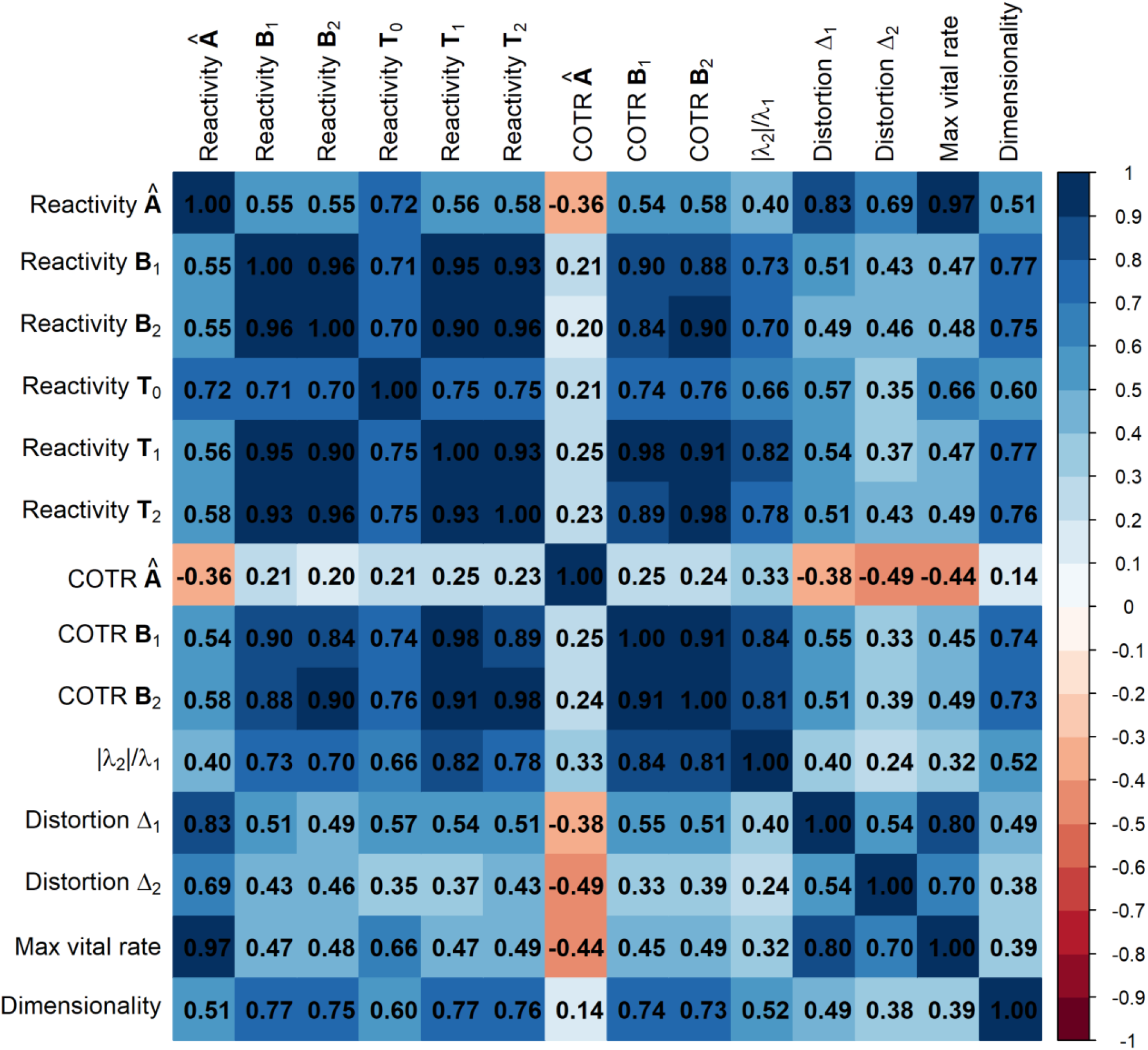
Spearman’s correlation matrix for 14 different measures: reactivities of 6 PPMs, COTRs of 3 PPMs, reciprocal of damping ratio, 2 different distortions, maximum vital rate, and dimensionality. The matrix norm used in the definition of reactivities and COTRs is the Frobenius norm.

### 3.2. Total responses and transient responses

Total response, measured by reactivity, was much smaller for balanced PPMs than for the original standardised PPMs (Figure 4a-c). For standardised PPMs 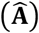, reactivities ranged from 1 to 593,753 (median = 2.97). For PPMs balanced with reproductive values (**B**_1_), reactivities ranged from 1 to 6.73 (median = 1.67). For PPMs balanced with stable stage densities (**B**_2_), reactivities ranged from 1 to 7.66 (median = 1.66). Spearman correlations between the reactivities of the standardised PPMs and those of the balanced matrices were moderate (*r*_*s*_ = 0.55 for both), whereas the Spearman correlation between the reactivities of the two balanced matrices was very strong (*r*_*s*_ = 0.96) (Figure 3).

**FIGURE 4.**
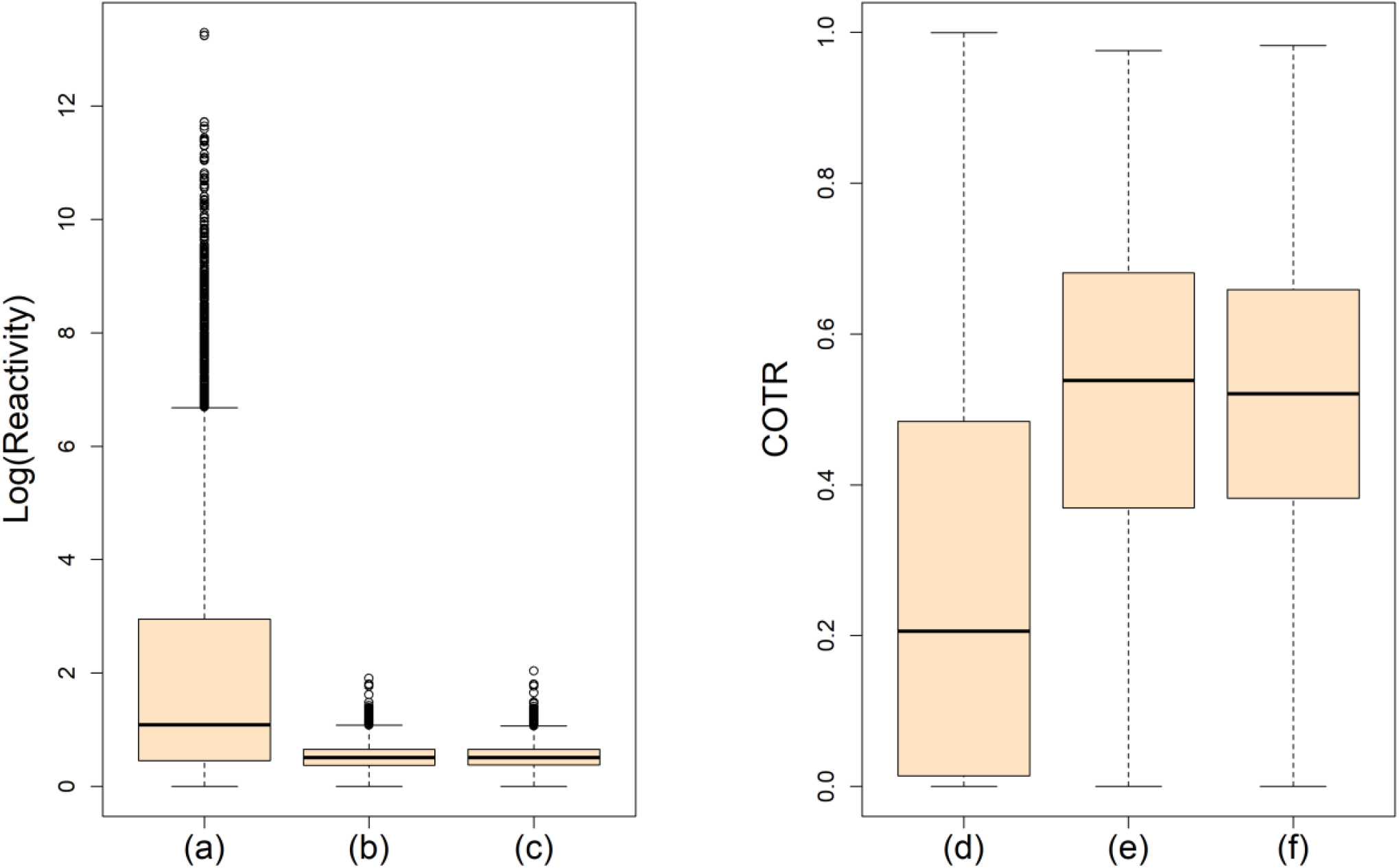
Box plots of distributions of reactivities (a)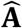, standardised PPMs; (b) **B**_1_, PPMs balanced with reproductive values; (c) **B**_2_, PPMs balanced with stable stage distribution, and COTRs applied to (d) 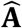, standardised PPMs; (e) **B**_1_, PPMs balanced with reproductive values; (f) **B**_2_, PPMs balanced with stable stage distribution. The matrix norm used in the definition of reactivities is the Frobenius norm. In each box plot, the horizontal line within the box indicates the median; the box encompasses 75% of the values, and the whiskers are drawn to the nearest value not beyond 1.5 × (interquartile range) from the quartiles.

Transient response, measured by COTR, tended to be smaller for standardised PPMs than for balanced matrices (Figure 4d-f). For the standardised PPMs 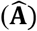, COTRs ranged from 0 to 1, (median = 0.21). For PPMs balanced with reproductive values (**B**_1_), COTRs ranged from 0 to 0.98 (median = 0.54). For PPMs balanced with stable stage distribution (**B**_2_), COTRs ranged from 0 to 0.98 (median = 0.52). Spearman correlations between the COTRs of the standardised PPMs and those of the balanced matrices were weak (*r*_*s*_ ≤ 0.25), whereas the Spearman correlation between the COTRS of the two balanced matrices was very strong (*r*_*s*_ = 0.91) (Figure 3).

I compared the distributions of standardised versus balanced PPMs using box plots of total response (reactivity) grouped by taxonomic order and ranked the taxonomic orders by median reactivity (Figure 5). There were large differences between the rankings of taxonomic orders when reactivities were applied to balanced rather than standardised PPMs. The mean absolute difference in ranks using 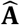 versus **B**_1_ and 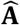 versus **B**_2_ were 6.9 and 5.8, respectively.

**FIGURE 5.**
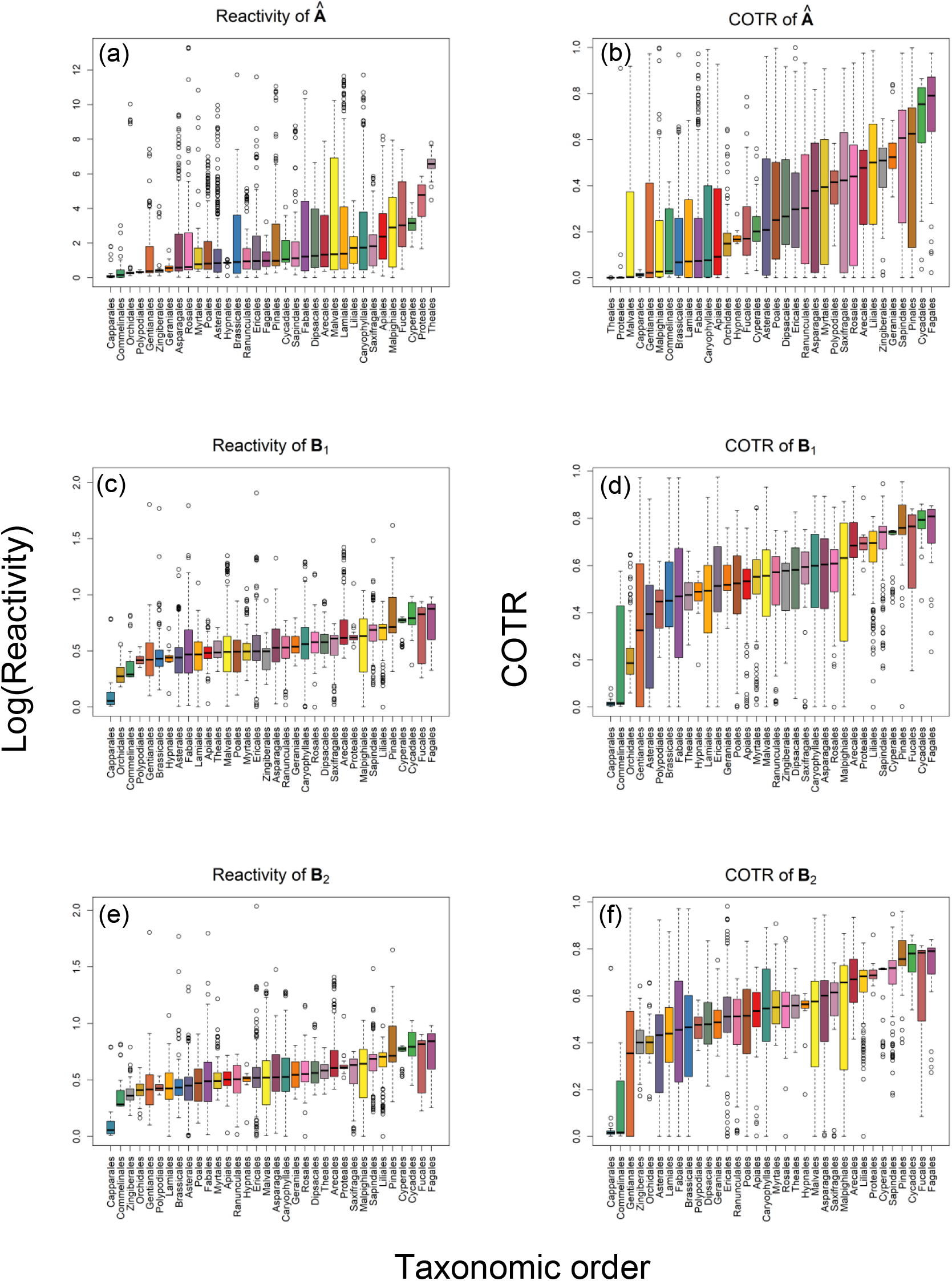
Box plots of reactivities representing total response and COTRs representing transient response. The top panels show (a) log(reactivity) and (b) COTR of 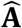, the standardised PPMs. The middle panels show (c) log(reactivity) and (d) COTR of **B**_1_, PPMs balanced with reproductive values. The bottom panels show (e) log(reactivity) and (f) COTR of **B**_2_, PPMs balanced with stable stage distribution. The matrix norm used in the definition of reactivities and COTRs is the Frobenius norm. The box plots, coloured by taxonomic order, are arranged so that the medians are increasing. In each box plot, the horizontal line within the box indicates the median; the box encompasses 75% of the values, and the whiskers are drawn to the nearest value not beyond 1.5 × (interquartile range) from the quartiles.

The rankings were relatively consistent between the two types of balanced PPMs. The mean absolute difference in ranks using **B**_1_ versus **B**_2_ was 2.1.

I also compared the distributions of the transient response, measured by COTR, of standardised versus balanced PPMs using box plots grouped by taxonomic order and ranked the taxonomic orders by median COTR (Figure 5). There were large differences between the rankings of taxonomic orders for balanced versus standardised PPMs. The mean absolute difference in ranks using 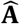 versus **B**_1_ and 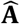 versus **B**_2_ were 6.8 and 7.8, respectively. The rankings were again relatively consistent between the two types of balanced PPMs. The mean absolute difference in ranks using **B**_1_ versus **B**_2_ was 2.8.

### 3.3. Reactivities vs. distortions

Distortion had a greater influence on the reactivities of the original standardised PPMs than they did on the reactivities of balanced PPMs and transient PPMs (Figure 3). Spearman correlations between the reactivity of 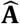 and the distortions Δ_1_ and Δ_2_ were strong: *r*_*s*_ = 0.83 and *r*_*s*_ = 0.69, respectively. In contrast, Spearman correlations showed that distortions were moderately correlated with the reactivities of the balanced matrices (0.43 ≤ *r*_*s*_ ≤ 0.51), and were weakly-to-moderately correlated with the reactivities of the transient matrices (0.35 ≤ *r*_*s*_ ≤ 0.57).

### 3.4. Transient dynamics vs. dimensionality

Previous studies found that transient dynamics were influenced by matrix dimensionality (Hinrichsen, 2023; Stott et al., 2010; Tenhumberg et al., 2009). I tested for the influence of dimensionality using the Spearman’s rank correlation matrix (Figure 3). Spearman correlations confirmed that dimensionality was highly correlated with the reactivities of **T**_1_, and **T**_2_ (*r*_*s*_ ≥ 0.76) with the COTRs of **B**_1_ and **B**_2_ (*r*_*s*_ ≥ 0.73).

### 3.5. Life histories that foster the greatest transient dynamics

Life histories that foster the greatest transient dynamics were discovered by finding PPMs with *ρ*(**T**_1_) and *ρ*(**T**_1_) equal to their upper bounds of 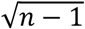. There were six such PPMs, all imprimitive. These PPMs were Leslie matrices for which only the oldest age class had a nonzero fertility rate. Moreover, **B**_1_ = **B**_2_ and **T**_1_ = **T**_2_. Their standardised PPMs had *n* eigenvalues of magnitude 1, a condition known to foster permanent waves (Bernadelli, 1941; Caswell, 2001).

## 4. DISCUSSION

This article presents a new analytical framework to measure transient dynamics that addresses three core challenges of the conventional framework: undue influence of stages that are numerous but of low value, scale dependence, and conflation of transient and asymptotic population dynamics. The first two challenges are remedied by balancing the PPM either by its reproductive values (i.e. Fisher balancing) or by its stable stage distribution (i.e. Demetrius balancing). The third challenge is solved by stripping the output of a PPM (balanced or not) of its component in the direction of its stable stage distribution. Stripping yields a transient PPM, which is used in place of the standardised PPM when computing indices of transient dynamics. I define a new index of transient dynamics, COTR, which is the fraction of the sum of squares of the total response that is explained by the transient response.

Several changes materialised when I used the new analytical framework in place of the conventional framework to analyse PPMs in COMPADRE. There were three changes in total response observed using the new framework: (1) the influence of distortion on the measures of total response decreased; (2) the estimates of total response shrank, with the largest reduced by several orders of magnitude; (3) the rankings of taxonomic orders by total response changed dramatically. These changes occur because balancing limits the size of vital rates, ensuring that none is greater than 1. This, in turn, reduces distortion by reducing the exceptionally high fertility rates that produce many immature individuals of low value. These individuals exert an undue influence, thus distorting the reactivities of standardised PPMs.

Using the new framework in place of the conventional one, where transient response was measured as COTR, produced two notable changes: (1) the transient response was substantially higher; and (2) the rankings of taxonomic orders by transient response changed dramatically.

These changes occur because, without balancing, the total response is often interpreted as mainly asymptotic, causing a reduction in the measured transient response. This is another consequence of the undue influence of abundant stages of low value.

Scale-invariant measures are useful because ecologists need robust measures for comparative demography, where there are infinitely many parameterisations of the same model. Results should not depend on whether seeds (or other stages) are counted by the hundredths, tenths, ones, hundreds, thousands, etc., or if stages are counted using a pre-reproductive vs. a post-reproductive census. With scale-invariant indices, it does not matter if a stage is counted differently; if alternative counts are proportional, the indices will not change. Among the scale-invariant statistics already available for comparative demography are the following: stable growth rate, elasticity of stable growth rate to vital rates, damping ratio, generation time, and Demetrius’s entropy (Caswell, 2001). With balancing, indices of transient dynamics can join this collection of valuable matrix statistics.

When the Frobenius norm was used, the PPMs that fostered a large transient response were imprimitive with an index of imprimitivity equal to *n*. These were Leslie matrices representing nonoverlapping generations, with only the oldest age class reproducing. That the transient dynamics are maximal for these imprimitive PPMs makes biological sense because they produce waves that never die out (Bernadelli, 1941; Caswell, 2001).

There are more indices of transient dynamics to consider within this new framework. A more varied set of indices of transient dynamics, which are applied to standardised PPMs, can also be applied directly to transient PPMs. Such indices include amplification envelope, maximum amplification, Kriess bound, etc. (Stott et al., 2011; Townley, 2007). Matrix norms in addition to the Frobenius norm should also be explored. The damping ratio deserves a prominent place among the indices of transient dynamics. It has three important properties: It is scale invariant, its reciprocal is a lower bound on the reactivity of transient PPMs, and it is highly correlated with other scale invariant indices of transient dynamics (Figure 3).

An important decision to make when evaluating transient dynamics is whether to use Fisher balancing (i.e. balancing by reproductive values) or Demetrius balancing (i.e. balancing by stable stage distribution). Fisher balancing values stages *prospectively*, while Demetrius balancing values stages *retrospectively*, which is revealed by how they value post-reproductive classes. When Fisher balancing is used, post-reproductive individuals have no value because they cannot contribute to future generations. However, when Demetrius balancing is used, post-reproductive individuals can be highly valued because, assuming a low survival probability from the youngest age class to maturity, many younger individuals were required to produce them. To meet conservation goals, Fisher balancing is perhaps most suitable; otherwise, a population that consists mainly of individuals of low reproductive potential may be overvalued, disguising a threat to population viability.

Studies of transient dynamics should consider the influence of dimensionality (Hinrichsen, 2023; Stott et al., 2010; Tenhumberg et al., 2009). To compare the transient dynamics of PPMs of different dimensionalities, one could collapse the larger PPM so that it has the same dimensionality as the smaller PPM (Hinrichsen, 2023; Hooley, 2000; Salguero-Gómez & Casper, 2010). Or, alternatively, one could account for dimensionality by including it as an explanatory variable in a multiple regression framework (Yokomizo et al., 2024).

An important future direction is to examine the sensitivity of the indices of transient dynamics to the vitality rates in the original PPM. Sensitivity analysis can help us understand what specific vital rates contribute most (or least) to transient dynamics (Caswell, 2007; Fox & Gurevitch, 2000; Yearsley, 2004). Another future direction is to adopt this new analytical framework to integral projection models (IPM), which use continuous stages (Doak et al., 2021; Merow et al., 2014). It is also natural to extend the analytical framework to stochastic matrices where vital rates vary over time according to probability distributions (Caswell, 2001).

## 5. CONCLUSIONS

The conventional framework of transient dynamics needs to be revised due to the three core challenges highlighted in this article. I present a revised framework that solves all three challenges by weighting stages by their values and disentangling asymptotic and transient dynamics. The revised framework leads to new inferences about the ranking of plant populations from smallest to largest transient dynamics. This finding suggests that previous comparative studies on transient dynamics should be re-evaluated using the revised framework. By solving the challenges with balancing and stripping, the revised framework offers a clearer and more robust portrait of transient dynamics for comparative demography.

## Acknowledgements

I gratefully acknowledge the COM(P)ADRE Team and its funding bodies for compiling and maintaining the demographic data that allowed me to conduct my analysis.

