## Appendix S1 for "Solving three core challenges in transient dynamics analysis of matrix population models"

### APPENDIX S1: BALANCING MAKES INDICES OF TRANSIENT DYNAMICS SCALE INVARIANT

In this appendix, I demonstrate that the balanced PPMs $\mathbf{B}_1$ and $\mathbf{B}_2$ and their transient forms $\mathbf{T}_1$ and $\mathbf{T}_2$, respectively, are scale invariant. Consequently, reactivity and COTR are also scale invariant when applied to these PPMs. Balancing also makes case-specific reactivities and COTRs scale-invariant. To show scale invariance, I rescale the model and demonstrate that, with balancing, the reactivities and COTRs derived from the rescaled model are identical to those derived from the original model. Scale invariance holds whether balancing is accomplished with reproductive values (Fisher balancing) or the stable stage distribution (Demetrius balancing).

Recall that the original model is governed by

$$\mathbf{n}_{t+1} = \mathbf{A}\mathbf{n}_t, \tag{S1.1}$$

where $\mathbf{n}_t$ is the stage vector at time $t$ with $n$ components, and $\mathbf{A}$ is an $n \times n$ PPM. Let $\mathbf{v}$ be the vector of reproductive values and let $\mathbf{w}$ be the stable stage distribution associated with dominant eigenvalue $\lambda_1$, such that $\mathbf{w}$ sums to 1 and $\mathbf{v}^\top\mathbf{w} = 1$. Balancing $\mathbf{A}$ with reproductive values yields the balanced PPM

$$\mathbf{B}_1 = \mathrm{diag}(\mathbf{v})\widehat{\mathbf{A}}\{\mathrm{diag}(\mathbf{v})\}^{-1}, \tag{S1.2}$$

where $\widehat{\mathbf{A}} = \mathbf{A}/\lambda_1$ is the standardized PPM. The PPM $\mathbf{B}_1$ has a dominant eigenvalue of 1 because it is similar to $\widehat{\mathbf{A}}$ via $\{\mathrm{diag}(\mathbf{v})\}^{-1}$, and therefore it has the same eigenvalues as $\widehat{\mathbf{A}}$ (Horn & Johnson, 2013).

Next consider a rescaled version of the original matrix population model where the stages of the rescaled model are defined by

$$\mathbf{n}'_t = \mathrm{diag}(\mathbf{c})\mathbf{n}_t, \tag{S1.3}$$

where $\mathbf{c}$ is an $n$-vector of positive weights. The equation governing the dynamics of $\mathbf{n}'_t$ is

$$\begin{aligned} \mathbf{n}'_{t+1} &= \mathrm{diag}(\mathbf{c})\mathbf{n}_{t+1} \\ &= \mathrm{diag}(\mathbf{c})\mathbf{A}\mathbf{n}_t \\ &= \mathrm{diag}(\mathbf{c})\mathbf{A}\{\mathrm{diag}(\mathbf{c})\}^{-1}\boldsymbol{n}'_t \\ &= \mathbf{A}'\,\mathbf{n}'_{t,} \end{aligned} \tag{S1.4}$$

where $\mathbf{A}' = \mathrm{diag}(\mathbf{c})\mathbf{A}\{\mathrm{diag}(\mathbf{c})\}^{-1}$ is the PPM for the rescaled model. I show that the balanced version of $\mathbf{A}'$, call it $\mathbf{B}'_1$, is equal to the balanced version of $\mathbf{A}$, namely $\mathbf{B}_1$, and therefore $\mathbf{B}_1$ is scale invariant. First, $\mathbf{A}$ and $\mathbf{A}'$ have the same eigenvalues because $\mathbf{A}$ is similar to $\mathbf{A}'$ via $\{\mathrm{diag}(\mathbf{c})\}^{-1}$ (Horn & Johnson, 2013). Furthermore, if $\mathbf{v}$ is the vector of reproductive values of $\mathbf{A}$ associated with $\lambda_1$, then $\mathbf{v}' = \mathbf{c}^\top\mathbf{w}\{\mathrm{diag}(\mathbf{c})\}^{-1}\mathbf{v}$ is the vector of reproductive values of $\mathbf{A}'$ and $\mathbf{w}' = \mathrm{diag}(\mathbf{c})\boldsymbol{w}/\mathbf{c}^\top\mathbf{w}$ is the stable stage distribution of $\mathbf{A}'$ associated with $\lambda_1$. As usual, the vectors $\mathbf{v}'$ and $\mathbf{w}'$ are scaled so that $\mathbf{w}'$ sums to 1 and $\mathbf{v}'^\top\mathbf{w}' = 1$.

The following shows that $\mathbf{v}' = \mathbf{c}^\top\mathbf{w}\{\mathrm{diag}(\mathbf{c})\}^{-1}\mathbf{v}$ is indeed the vector of reproductive values of $\mathbf{A}'$:

$$\begin{aligned}
\mathbf{v}'^{\top}\mathbf{A}' &= (\mathbf{c}^{\top}\mathbf{w})\mathbf{v}^{\top}\{\mathrm{diag}(\mathbf{c})\}^{-1}\mathbf{A}' \\
&= (\mathbf{c}^{\top}\mathbf{w})\mathbf{v}^{\top}\{\mathrm{diag}(\mathbf{c})\}^{-1}\mathrm{diag}(\mathbf{c})\mathbf{A}\{\mathrm{diag}(\mathbf{c})\}^{-1} \\
&= (\mathbf{c}^{\top}\mathbf{w})\mathbf{v}^{\top}\mathbf{A}\{\mathrm{diag}(\mathbf{c})\}^{-1} \\
&= (\mathbf{c}^{\top}\mathbf{w})\lambda_1\mathbf{v}^{\top}\{\mathrm{diag}(\mathbf{c})\}^{-1} \\
&= \lambda_1\mathbf{v}'^{\top}.
\end{aligned} \tag{S1.5}$$

A similar argument shows that $\mathbf{w}' = \mathrm{diag}(\mathbf{c})\mathbf{w}/\mathbf{c}^{\top}\mathbf{w}$ is the stable stage distribution of $\mathbf{A}'$.

Armed with these findings, balancing $\mathbf{A}'$ with its reproductive values yields

$$\begin{aligned}
\mathbf{B}_1' &= \mathrm{diag}(\mathbf{v}')(\mathbf{A}'/\lambda_1)\{\mathrm{diag}(\mathbf{v}')\}^{-1} \\
&= (\mathbf{c}^{\top}\mathbf{w})\mathrm{diag}(\{\mathrm{diag}(\mathbf{c})\}^{-1}\mathbf{v})(\mathbf{A}'/\lambda_1)[\mathrm{diag}(\{\mathrm{diag}(\mathbf{c})\}^{-1}\mathbf{v})]^{-1}/(\mathbf{c}^{\top}\mathbf{w}) \\
&= \mathrm{diag}(\mathbf{v})\{\mathrm{diag}(\mathbf{c})\}^{-1}(\mathrm{diag}(\mathbf{c})\mathbf{A}\{\mathrm{diag}(\mathbf{c})\}^{-1}/\lambda_1)\mathrm{diag}(\mathbf{c})\{\mathrm{diag}(\mathbf{v})\}^{-1} \\
&= \mathrm{diag}(\mathbf{v})(\mathbf{A}/\lambda_1)\{\mathrm{diag}(\mathbf{v})\}^{-1} \\
&= \mathbf{B}_1.
\end{aligned} \tag{S1.6}$$

In brief, the weights **c** cancel out in the calculation of $\mathbf{B}_1'$, yielding the matrix $\mathbf{B}_1$. Therefore, $\mathbf{B}_1$ is scale invariant since it is unchanged by rescaling the original matrix population model. This, in turn, implies that the transient form $\mathbf{T}_1$ is scale-invariant, since it is completely determined by $\mathbf{B}_1$. Therefore, indices of transient dynamics, or any statistic for that matter, are scale invariant when they are applied to $\mathbf{B}_1$ or $\mathbf{T}_1$. A similar argument shows that indices of transient dynamics are scale invariant when they are applied to $\mathbf{B}_2$ or $\mathbf{T}_2$. Thus, the reactivities $\rho(\mathbf{B}_1)$, $\rho(\mathbf{B}_2)$, $\rho_1 = \rho(\mathbf{T}_1)$, and $\rho_2 = \rho(\mathbf{T}_2)$ are all scale invariant, regardless of the matrix norm used in the definition of reactivity. Furthermore, the COTRs $q_1^2$ and $q_2^2$, which are functions of these scale-invariant matrices, are also scale invariant.

The case-specific reactivities $\bar{\rho}_1$ and $\bar{\rho}_2$ are also scale invariant. Let us first demonstrate scale invariance for

$$\bar{\rho}_1 = \frac{\|\mathbf{T}_1 \mathbf{V} \mathbf{n}_0\|}{\|\mathbf{V} \mathbf{n}_0\|}, \tag{S1.7}$$

where $\|\cdot\|$ is any vector norm, and $\mathbf{V} = \mathrm{diag}(\mathbf{v})$. To show that $\bar{\rho}_1$ is scale-invariant, we rescale the original model as in Equation (S1.3). The versions of $\mathbf{n}_0$, $\mathbf{v}$, and $\mathbf{T}_1$ for the rescaled model are $\mathbf{n}_0' = \mathrm{diag}(\mathbf{c})\mathbf{n}_0$, $\mathbf{v}' = \mathbf{c}^\top \mathbf{w}\{\mathrm{diag}(\mathbf{c})\}^{-1}\mathbf{v}$ and $\mathbf{T}_1' = \mathbf{T}_1$, respectively. The version of Equation (S1.7) for the rescaled model is

$$\begin{aligned}
\bar{\rho}_1' &= \frac{\|\mathbf{T}_1' \mathrm{diag}(\mathbf{v}')\mathbf{n}_0'\|}{\|\mathrm{diag}(\mathbf{v}')\mathbf{n}_0'\|} \\
&= \frac{\|\mathbf{T}_1 \mathbf{c}^\top \mathbf{w}\{\mathrm{diag}(\mathbf{c})\}^{-1}\mathrm{diag}(\mathbf{v})\mathrm{diag}(\mathbf{c})\mathbf{n}_0\|}{\|\mathbf{c}^\top \mathbf{w}\{\mathrm{diag}(\mathbf{c})\}^{-1}\mathrm{diag}(\mathbf{v})\mathrm{diag}(\mathbf{c})\mathbf{n}_0\|} \\
&= \frac{\mathbf{c}^\top \mathbf{w}\|\mathbf{T}_1\{\mathrm{diag}(\mathbf{c})\}^{-1}\mathrm{diag}(\mathbf{v})\mathrm{diag}(\mathbf{c})\mathbf{n}_0\|}{\mathbf{c}^\top \mathbf{w}\|\{\mathrm{diag}(\mathbf{c})\}^{-1}\mathrm{diag}(\mathbf{v})\mathrm{diag}(\mathbf{c})\mathbf{n}_0\|} \\
&= \frac{\|\mathbf{T}_1 \mathrm{diag}(\mathbf{v})\{\mathrm{diag}(\mathbf{c})\}^{-1}\mathrm{diag}(\mathbf{c})\mathbf{n}_0\|}{\|\mathrm{diag}(\mathbf{v})\{\mathrm{diag}(\mathbf{c})\}^{-1}\mathrm{diag}(\mathbf{c})\mathbf{n}_0\|} \\
&= \frac{\|\mathbf{T}_1 \mathrm{diag}(\mathbf{v})\mathbf{n}_0\|}{\|\mathrm{diag}(\mathbf{v})\mathbf{n}_0\|} \\
&= \bar{\rho}_{1.}
\end{aligned} \tag{S1.8}$$

The second line expresses quantities derived from the rescaled model in terms of their corresponding quantities from the original model. The third line follows because vector norms are homogenous, which allows us to bring the scalar $\mathbf{c}^\top \mathbf{w}$ outside of the norms in the numerator and denominator (Horn & Johnson, 2013). Line four follows because diagonal matrices commute. Therefore, $\bar{\rho}_1' = \bar{\rho}_1$. Similarly, we can show that $\bar{\rho}_2' = \bar{\rho}_2$. Thus, with balancing, case-specific reactivities are scale-invariant.

Lastly, I show that case-specific COTRs $\bar{q}_1^2$ and $\bar{q}_2^2$ are also scale invariant. Let us first demonstrate scale invariance for

$$\bar{q}_1^2 = \frac{\|\mathbf{T}_1\mathbf{V}\mathbf{n}_0\|_2^2}{\|\mathbf{B}_1\mathbf{V}\mathbf{n}_0\|_2^2}, \tag{S1.9}$$

where $\|\cdot\|_2$ is the Euclidean vector norm. To show that $\bar{q}_1^2$ is scale-invariant, we rescale the original model as in Equation (S1.3). The versions of $\mathbf{n}_0$, $\mathbf{v}$, $\mathbf{T}_1$, and $\mathbf{B_1}$ for the rescaled model are of $\mathbf{n}_0' = \mathrm{diag}(\mathbf{c})\mathbf{n}_0$, $\mathbf{v}' = \mathbf{c}^\top\mathbf{w}\{\mathrm{diag}(\mathbf{c})\}^{-1}\mathbf{v}$, $\mathbf{T}_1' = \mathbf{T}_1$, and $\mathbf{B}_1' = \mathbf{B_1}$. The version of Equation (S1.9) for the rescaled model is

$$\begin{aligned}
\bar{q}_1'^2 &= \frac{\|\mathbf{T}_1'\mathrm{diag}(\mathbf{v}')\mathbf{n}_0'\|_2^2}{\|\mathbf{B}_1'\mathrm{diag}(\mathbf{v}')\mathbf{n}_0'\|_2^2} \\
&= \frac{\|\mathbf{T}_1\mathbf{c}^\top\mathbf{w}\{\mathrm{diag}(\mathbf{c})\}^{-1}\mathrm{diag}(\mathbf{v})\mathrm{diag}(\mathbf{c})\mathbf{n}_0\|_2^2}{\|\mathbf{B}_1\mathbf{c}^\top\mathbf{w}\{\mathrm{diag}(\mathbf{c})\}^{-1}\mathrm{diag}(\mathbf{v})\mathrm{diag}(\mathbf{c})\mathbf{n}_0\|_2^2} \\
&= \frac{(\mathbf{c}^\top\mathbf{w})^2\|\mathbf{T}_1\{\mathrm{diag}(\mathbf{c})\}^{-1}\mathrm{diag}(\mathbf{v})\mathrm{diag}(\mathbf{c})\mathbf{n}_0\|_2^2}{(\mathbf{c}^\top\mathbf{w})^2\|\mathbf{B}_1\{\mathrm{diag}(\mathbf{c})\}^{-1}\mathrm{diag}(\mathbf{v})\mathrm{diag}(\mathbf{c})\mathbf{n}_0\|_2^2} \\
&= \frac{\|\mathbf{T}_1\mathrm{diag}(\mathbf{v})\{\mathrm{diag}(\mathbf{c})\}^{-1}\mathrm{diag}(\mathbf{c})\mathbf{n}_0\|_2^2}{\|\mathbf{B}_1\mathrm{diag}(\mathbf{v})\{\mathrm{diag}(\mathbf{c})\}^{-1}\mathrm{diag}(\mathbf{c})\mathbf{n}_0\|_2^2} \\
&= \frac{\|\mathbf{T}_1\mathrm{diag}(\mathbf{v})\mathbf{n}_0\|_2^2}{\|\mathbf{B}_1\mathrm{diag}(\mathbf{v})\mathbf{n}_0\|_2^2} \\
&= \bar{q}_1^2.
\end{aligned} \tag{S1.10}$$

The second line expresses quantities derived from the rescaled model in terms of their corresponding quantities from the original model. The third line follows because vector norms are homogenous, which allows us to bring the scalar $\mathbf{c}^\top\mathbf{w}$ outside of the norms in the numerator and denominator. Line four follows because diagonal matrices commute. Therefore, $\bar{q}_1'^2 = \bar{q}_1^2$. Similarly, we can show that $\bar{q}_2'^2 = \bar{q}_2^2$. Thus, with balancing, the case-specific COTRs are scale-invariant.
