## Appendix S2 for "Solving three core challenges in transient dynamics analysis of matrix population models"

**APPENDIX S2: BOUNDS ON REACTIVITIES**

In this appendix, I derive bounds on reactivities for PPMs $\widehat{\mathbf{A}}$, $\mathbf{B}_1$,and $\mathbf{B}_2$,; their transient forms $\mathbf{T}_0$,$\mathbf{T}_1$, and $\mathbf{T}_2$; and their asymptotic forms, $\mathbf{Q}_0$,$\mathbf{Q}_1$, and $\mathbf{Q}_2$. Because they were derived without balancing, only lower bounds will be supplied for reactivities of $\widehat{\mathbf{A}}$, $\mathbf{T}_0$, and $\mathbf{Q}_0$. To derive lower bounds on the reactivities, I use the fact that the spectral radius is a lower bound for any matrix norm (Horn & Johnson, 2013). Throughout, reactivities will be defined using the Frobenius norm. I will show that the bounds on the balanced matrices are given by

$$1 \leq \|\mathbf{B}_k\|_2 \leq \sqrt{n} \qquad \text{(S2.1)}$$

where $k = 1{,}2$; and $n$ is the dimensionality of $\mathbf{A}$. Furthermore, $1 \leq \left\|\widehat{\mathbf{A}}\right\|_2$.

When $n > 1$ the bounds on the transient matrices are

$$\frac{|\lambda_2|}{\lambda_1} \leq \|\mathbf{T}_k\|_2 \leq \sqrt{n-1} \qquad \text{(S2.2)}$$

where $|\lambda_2|$ is the magnitude of the second largest eigenvalue of $\mathbf{A}$, and $\lambda_1$ is the largest eigenvalue of $\mathbf{A}$, and $k = 1{,}2$. Furthermore, $\frac{|\lambda_2|}{\lambda_1} \leq \|\mathbf{T}_0\|_2$. When $n = 1$, $\|\mathbf{T}_k\|_2 = 0$ ($k = 0{,}1{,}2$).

The bounds on the asymptotic PPMs are

$$1 \leq \|\mathbf{Q}_k\|_2 \leq \sqrt{n} \qquad \text{(S2.3)}$$

where $k = 1{,}2$. Furthermore, $1 \leq \|\mathbf{Q}_0\|_2$.

When matrix norms besides the Frobenius norm are used, the lower bounds developed here will continue to apply because the lower bounds are built on the spectral radiuses of the PPMs, and the spectral radius is a lower bound regardless of the matrix norm used (Horn & Johnson, 2013).

**Bounds for the reactivities of $\widehat{\mathbf{A}}$, $\mathbf{B}_1$, and $\mathbf{B}_2$**

The lower bound on reactivity of these three PPMs is 1 because a matrix norm is always greater than or equal to the magnitude of the largest eigenvalue. Therefore, $\rho(\widehat{\mathbf{A}}) \geq 1$, $\rho(\mathbf{B}_1) \geq 1$, and $\rho(\mathbf{B}_2) \geq 1$.

Next, I prove that the upper bound on reactivity of the transient matrix, $\mathbf{B}_1$, is $\sqrt{n}$. First let $\mathbf{b}$ be any column $\mathbf{B}_1$ with the largest Euclidean norm. I will show that $||\mathbf{b}||_2 \leq 1$, and since $||\mathbf{B}_1||_2^2$ is the sum of the norm squared of all columns, then $||\mathbf{B}_1||_2 \leq \sqrt{n}$. Let $\mathbf{e}_i$ represent the standard basis vectors, which are the columns of the identity matrix. Then

$$\begin{aligned} ||\mathbf{b}||_2 &= \left\|\sum_{i=1}^{n} \mathrm{b}_i \mathbf{e}_i\right\|_2 \\ &\leq \sum_{i=1}^{n} ||\mathrm{b}_i \mathbf{e}_i||_2 \\ &\leq \sum_{i=1}^{n} ||\mathrm{b}_i||_2 \, ||\mathbf{e}_i||_2 \\ &= \sum_{i=1}^{n} |\mathrm{b}_i| \\ &= 1 \end{aligned} \tag{S2.4}$$

The second line follows from the triangle inequality, the third line follows by the Cauchy–Schwarz inequality, the fourth line follows since $||\mathbf{e}_i||_2 = 1$, and the last follows since the columns of $\mathbf{B}_1$ sum to 1. Therefore,

$$\rho^2(\mathbf{B}_1) = \|\mathbf{B}_1\|_2^2 \qquad \text{(S2.5)}$$
$$\leq n\|\mathbf{b}\|_2^2$$
$$\leq n$$

Thus, $1 \leq \rho(\mathbf{B}_1) \leq \sqrt{n}$. A similar argument shows that $1 \leq \rho(\mathbf{B}_2) \leq \sqrt{n}$.

**Bounds on the reactivities of $\mathbf{T}_0$ $\mathbf{T}_1$ and $\mathbf{T}_2$**

When $n > 1$, the lower bound on any transient matrix is $\frac{|\lambda_2|}{\lambda_1}$ since any norm of a matrix is greater than or equal to the spectral radius of the matrix, and the spectral radius of the transient PPMs is $\frac{|\lambda_2|}{\lambda_1}$. Therefore, $\rho(\mathbf{T}_k) \geq \frac{|\lambda_2|}{\lambda_1}$ for $(k = 0,1,2)$. I next show that the upper bound of $\rho(\mathbf{T}_1)$ is $\sqrt{n-1}$. Using the fact that $\|\mathbf{B}_1\|_2^2 = \|\mathbf{T}_1\|_2^2 + \|\mathbf{Q}_1\|_2^2$ which I demonstrate in Appendix S3,

$$\rho^2(\mathbf{T}_1) = \|\mathbf{T}_1\|_2^2 \qquad \text{(S2.6)}$$
$$= \|\mathbf{B}_1\|_2^2 - \|\mathbf{Q}_1\|_2^2$$
$$= \|\mathbf{B}_1\|_2^2 - \|\mathbf{P}_1\mathbf{B}_1\|_2^2$$
$$\leq n-1$$

The last line follows because $\|\mathbf{B}_1\|_2^2 \leq n$ and the largest eigenvalue of $\mathbf{P}_1\mathbf{B}_1$ is equal to 1, corresponding to right eigenvector $\mathbf{v} \circ \mathbf{w}$, and therefore $1 \leq \|\mathbf{P}_1\mathbf{B}_1\|_2$, because the norm of a matrix is always greater than or equal to the spectral radius of the matrix (Horn & Johnson, 2013). Therefore, when $n > 1$, $\frac{|\lambda_2|}{\lambda_1} \leq \rho(\mathbf{T}_1) \leq \sqrt{n-1}$. A similar argument shows that when $n > 1$, $\frac{|\lambda_2|}{\lambda_1} \leq \rho(\mathbf{T}_2) \leq \sqrt{n-1}$. When $n = 1$, then $\mathbf{P}_k = 1$ $(k = 0,1,2)$, and therefore $\mathbf{T}_0 = (1 - \mathbf{P}_0)\widehat{\mathbf{A}} = 0$, $\mathbf{T}_1 = (1 - \mathbf{P}_1)\mathbf{B}_1 = 0$, and $\mathbf{T}_2 = (1 - \mathbf{P}_2)\mathbf{B}_2 = 0$.

**Bounds on the reactivities of $\mathbf{Q}_0$ $\mathbf{Q}_1$ and $\mathbf{Q}_2$**

The lower bound on any asymptotic matrix is 1 since any norm of a matrix is greater than or equal to the spectral radius of the matrix, and the spectral radius of an asymptotic PPM is 1. Therefore, $\rho(\mathbf{Q}_k) \geq 1$ for $(k = 0,1,2)$. I next show that the upper bound of $\rho(\mathbf{Q}_1)$ is $\sqrt{n}$. Using the fact that $\|\mathbf{B}_1\|_2^2 = \|\mathbf{T}_1\|_2^2 + \|\mathbf{Q}_1\|_2^2$ which I demonstrate in Appendix S3,

$$\begin{aligned} \rho^2(\mathbf{Q}_1) &= \|\mathbf{Q}_1\|_2^2 \\ &= \|\mathbf{B}_1\|_2^2 - \|\mathbf{T}_1\|_2^2 \\ &\leq n. \end{aligned} \quad \text{(S2.7)}$$

The last line follows because $\|\mathbf{B}_1\|_2^2 \leq n$ and $\|\mathbf{T}_1\|_2^2 \geq 0$. Thus $1 \leq \rho(\mathbf{Q}_1) \leq \sqrt{n}$. A similar argument shows that $1 \leq \rho(\mathbf{Q}_2) \leq \sqrt{n}$.
