## Appendix S3 for "Solving three core challenges in transient dynamics analysis of matrix population models"

### APPENDIX S3: COEFFICIENT OF TRANSIENT RESPONSE (COTR)

In this appendix, I show that the transient and asymptotic matrices, used to define the coefficient of transient response are orthogonal and therefore, $\|\widehat{\mathbf{A}}\|_2^2 = \|\mathbf{T}_0\|_2^2 + \|\mathbf{Q}_0\|_2^2$, $\|\,\mathbf{B}_1\|_2^2 = \|\mathbf{T}_1\|_2^2 + \|\mathbf{Q}_1\|_2^2$, and $\|\,\mathbf{B}_2\|_2^2 = \|\mathbf{T}_2\|_2^2 + \|\mathbf{Q}_2\|_2^2$. It is enough to demonstrate the result for $\widehat{\mathbf{A}}$, then by identical arguments it will follow for $\mathbf{B}_1$ and $\mathbf{B}_2$. These results are used to show that coefficients of transient dynamics $q_0^2 = \frac{\|\mathbf{T}_0\|_2^2}{\|\widehat{\mathbf{A}}\|_2^2}, q_1^2 = \frac{\|\mathbf{T}_1\|_2^2}{\|\mathbf{B}_1\|_2^2}$, and $q_2^2 = \frac{\|\mathbf{T}_2\|_2^2}{\|\mathbf{B}_2\|_2^2}$ are less than or equal to 1. When COTR $q_k^2$ $(k = 0,1,2)$ is equal to zero, the response is purely asymptotic, and when $q_k^2$ is equal to 1, the response is purely transient. There is always some measure of asymptotic response. Therefore, $q_k^2$ can approach 1, but not equal to 1. I then show analogous bounds for the case-specific COTRs.

First, I write $\widehat{\mathbf{A}} = \mathbf{T}_0 + \mathbf{Q}_0$, which follows since, by definition, the asymptotic matrix is $\mathbf{Q}_0 = \widehat{\mathbf{A}} - \mathbf{T}_0$. Recall that the transient matrix is defined as $\mathbf{T}_0 = (\mathbf{I} - \mathbf{P}_0)\widehat{\mathbf{A}}$, where $\mathbf{P}_0 = \frac{\mathbf{w}\mathbf{w}^\top}{\mathbf{w}^\top\mathbf{w}}$ and $\mathbf{w}$ is the stable stage distribution of $\widehat{\mathbf{A}}$. To show orthogonality of the columns of $\mathbf{Q}_0$ with the columns of $\mathbf{T}_0$, it is enough to show that $\mathbf{Q}_0^\top\mathbf{T}_0 = \mathbf{0}$ (using the definition of orthogonality).

$$\begin{aligned}
\mathbf{Q}_0^\top\mathbf{T}_0 &= \left(\mathbf{P}_0\widehat{\mathbf{A}}\right)^\top(\mathbf{I} - \mathbf{P}_0)\widehat{\mathbf{A}} \\
&= \widehat{\mathbf{A}}^\top\,\mathbf{P}_0^\top(\mathbf{I} - \mathbf{P}_0)\widehat{\mathbf{A}} \\
&= \widehat{\mathbf{A}}^\top\,\mathbf{P}_0(\mathbf{I} - \mathbf{P}_0)\widehat{\mathbf{A}} \\
&= \widehat{\mathbf{A}}^\top\,(\mathbf{P}_0 - \mathbf{P}_0^2)\widehat{\mathbf{A}} \\
&= \widehat{\mathbf{A}}^\top\,(\mathbf{P}_0 - \mathbf{P}_0)\widehat{\mathbf{A}} \\
&= \mathbf{0}.
\end{aligned} \tag{S3.1}$$

The third line of Equation (S3.1) follows because $\mathbf{P}_0^\top = \left(\frac{\mathbf{w}\mathbf{w}^\top}{\mathbf{w}^\top\mathbf{w}}\right)^\top = \frac{\mathbf{w}\mathbf{w}^\top}{\mathbf{w}^\top\mathbf{w}} = \mathbf{P}_0$. The fifth line follows because $\mathbf{P}_0 = \mathbf{P}_0^2$. Therefore, the columns of $\mathbf{P}_0$ are orthogonal to the columns of $\mathbf{Q}_0$. Next, I show that orthogonality implies that $\left\|\widehat{\mathbf{A}}\right\|_2^2 = \|\mathbf{T}_0\|_2^2 + \|\mathbf{Q}_0\|_2^2$.

$$\begin{aligned}
\left\|\widehat{\mathbf{A}}\right\|_2^2 &= \|\mathbf{T}_0 + \mathbf{Q}_0\|_2^2 \\
&= \mathrm{tr}\{(\mathbf{T}_0 + \mathbf{Q}_0)^\top(\mathbf{T}_0 + \mathbf{Q}_0)\} \\
&= \mathrm{tr}(\mathbf{T}_0^\top\mathbf{T}_0 + \mathbf{T}_0^\top\mathbf{Q}_0 + \mathbf{Q}_0^\top\mathbf{T}_0 + \mathbf{Q}_0^\top\mathbf{Q}_0) \\
&= \mathrm{tr}(\mathbf{T}_0^\top\mathbf{T}_0 + \mathbf{Q}_0^\top\mathbf{Q}_0) \\
&= \mathrm{tr}(\mathbf{T}_0^\top\mathbf{T}_0) + \mathrm{tr}(\mathbf{Q}_0^\top\mathbf{Q}_0) \\
&= \|\mathbf{T}_0\|_2^2 + \|\mathbf{Q}_0\|_2^2,
\end{aligned} \tag{S3.2}$$

where $\mathrm{tr}(\cdot)$ represents the trace of a matrix, which is the sum of its diagonal elements. The fourth line of Equation (S3.2) follows because, as we showed in Equation (S3.1), $\mathbf{Q}_0^\top\mathbf{T}_0 = \mathbf{0}$, and therefore, $\mathbf{T}_0^\top\mathbf{Q}_0 = (\mathbf{Q}_0^\top\mathbf{T}_0)^\top = \mathbf{0}$. Thus, Equation (S3.2) demonstrates that $\left\|\widehat{\mathbf{A}}\right\|_2^2 = \|\mathbf{T}_0\|_2^2 + \|\mathbf{Q}_0\|_2^2$. Identical arguments show that $\|\mathbf{B}_1\|_2^2 = \|\mathbf{T}_1\|_2^2 + \|\mathbf{Q}_1\|_2^2$, and $\|\mathbf{B}_2\|_2^2 = \|\mathbf{T}_2\|_2^2 + \|\mathbf{Q}_2\|_2^2$.

From Equation (S3.2), it immediately follows that

$$q_0^2 = \frac{\|\mathbf{T}_0\|_2^2}{\left\|\widehat{\mathbf{A}}\right\|_2^2}, \tag{S3.3}$$

must be less than or equal to 1. Identical arguments show that $q_1^2 = \frac{\|\mathbf{T}_1\|_2^2}{\|\mathbf{B}_1\|_2^2}$, and $q_2^2 = \frac{\|\mathbf{T}_2\|_2^2}{\|\mathbf{B}_2\|_2^2}$ are less than or equal to 1. Furthermore, the COTR $q_k^2$ $(k = 0,1,2)$ is equal to zero whenever $\mathbf{T}_k = \mathbf{0}$ (response is purely asymptotic) and equal to 1 whenever $\mathbf{Q}_k = \mathbf{0}$ (response is purely transient).

Note that $\mathbf{Q}_k = \mathbf{0}$ is not actually possible, because $\|\mathbf{Q}_k\|_2 \geq 1$ (Appendix S2). Therefore, in reality, $q_k^2$ can get close to 1, but not equal 1. There will always be some measure of asymptotic response.

I next show the analogous results for the case-specific versions of the previous equations. Let $\mathbf{n}_0$ be any nonzero initial stage vector used in the original matrix population model of Equation (1) of the main text. Then

$$\begin{aligned}\left\|\widehat{\mathbf{A}}\mathbf{n}_0\right\|_2^2 &= \|\mathbf{T}_0\mathbf{n}_0 + \mathbf{Q}_0\mathbf{n}_0\|_2^2 \\ &= (\mathbf{T}_0\mathbf{n}_0 + \mathbf{Q}_0\mathbf{n}_0)^\top(\mathbf{T}_0\mathbf{n}_0 + \mathbf{Q}_0\mathbf{n}_0) \\ &= \mathbf{n}_0^\top\mathbf{T}_0^\top\mathbf{T}_0\mathbf{n}_0 + \mathbf{n}_0^\top\mathbf{T}_0^\top\mathbf{Q}_0\mathbf{n}_0 + \mathbf{n}_0^\top\mathbf{Q}_0^\top\mathbf{T}_0\mathbf{n}_0 + \mathbf{n}_0^\top\mathbf{Q}_0^\top\mathbf{Q}_0\mathbf{n}_0 \\ &= \mathbf{n}_0^\top\mathbf{T}_0^\top\mathbf{T}_0\mathbf{n}_0 + \mathbf{n}_0^\top\mathbf{Q}_0^\top\mathbf{Q}_0\mathbf{n}_0 \\ &= \|\mathbf{T}_0\mathbf{n}_0\|_2^2 + \|\mathbf{Q}_0\mathbf{n}_0\|_2^2,\end{aligned} \tag{S3.4}$$

where $\|\cdot\|_2$ is the Euclidean vector norm. The fourth line of Equation (S3.4) follows because, as we showed in Equation (S3.1), $\mathbf{Q}_0^\top\mathbf{T}_0 = \mathbf{0}$. Thus, Equation (S3.4) demonstrates that $\left\|\widehat{\mathbf{A}}\mathbf{n}_0\right\|_2^2 = \|\mathbf{T}_0\mathbf{n}_0\|_2^2 + \|\mathbf{Q}_0\mathbf{n}_0\|_2^2$. Identical arguments show that $\|\mathbf{B}_1\mathbf{V}\mathbf{n}_0\|_2^2 = \|\mathbf{T}_1\mathbf{V}\mathbf{n}_0\|_2^2 + \|\mathbf{Q}_1\mathbf{V}\mathbf{n}_0\|_2^2$, and $\left\|\mathbf{B}_2\mathbf{W}^{-1}\mathbf{n}_0\right\|_2^2 = \left\|\mathbf{T}_2\mathbf{W}^{-1}\mathbf{n}_0\right\|_2^2 + \left\|\mathbf{Q}_2\mathbf{W}^{-1}\mathbf{n}_0\right\|_2^2$.

From Equation (S3.4), it follows that

$$\bar{q}_0^2 = \frac{\|\mathbf{T}_0\mathbf{n}_0\|_2^2}{\left\|\widehat{\mathbf{A}}\mathbf{n}_0\right\|_2^2} \tag{S3.5}$$

must be less than or equal to 1. Identical arguments show that $\bar{q}_1^2 = \frac{\|\mathbf{T}_1\mathbf{V}\mathbf{n}_0\|_2^2}{\|\mathbf{B}_1\mathbf{V}\mathbf{n}_0\|_2^2}$, and $\bar{q}_2^2 = \frac{\|\mathbf{T}_2\mathbf{W}^{-1}\mathbf{n}_0\|_2^2}{\|\mathbf{B}_2\mathbf{W}^{-1}\mathbf{n}_0\|_2^2}$ are less than or equal to 1. When $\bar{q}_k^2 = 0$ $(k = 0,1,2)$, the transient response is zero (response is purely asymptotic). When $\bar{q}_k^2 = 1$, the transient response is at a maximum (response is purely transient). In reality, $\bar{q}_k^2$ can get close to 1, but not equal 1, because there will always be a nonzero component of $\mathbf{n}_0$ in the direction of the stable stage distribution.
